# Boundary domain genes were recruited to suppress bract growth and promote branching in maize

**DOI:** 10.1101/2021.10.05.463134

**Authors:** Yuguo Xiao, Jinyan Guo, Zhaobin Dong, Annis Richardson, Erin Patterson, Sidney Mangrum, Seth Bybee, Edoardo Bertolini, Madelaine Bartlett, George Chuck, Andrea L. Eveland, Michael J. Scanlon, Clinton Whipple

## Abstract

Grass inflorescence development is diverse and complex and involves sophisticated but poorly understood interactions of genes regulating branch determinacy and leaf growth. Here, we use a combination of transcript profiling, genetic and phylogenetic analyses to investigate *tasselsheath1* (*tsh1*) and *tsh4*, two maize genes that simultaneously suppress inflorescence leaf growth and inhibit branching. We identify a regulatory network of inflorescence leaf suppression that involves the phase change gene *tsh4* upstream of *tsh1* and the ligule identity gene *liguleless2* (*lg2*). We also find that a series of duplications in the *tsh1* gene lineage facilitated its shift from boundary domain in non-grasses to suppressed inflorescence leaves of grasses. Collectively, these results suggest that the boundary domain genes *tsh1* and *lg2* were recruited to inflorescence leaves where they suppress growth and regulate a non-autonomous signaling center that promotes inflorescence branching, an important component of yield in cereal grasses.

## Introduction

The transition from vegetative to reproductive growth in plants is typically accompanied by dramatic morphological changes. Among these changes, leaf outgrowth, the dominant vegetative characteristic in most plants, is often highly reduced or completely suppressed. Leaves subtending reproductive structures (inflorescence branches or flowers) are called bracts, and some level of bract reduction or suppression is common, but not universal in the angiosperms (*1*).

Similar to most grasses, maize suppresses a subset of inflorescence bracts, which are only visible as a small ridge during early stages of inflorescence development (*2, 3*). While some inflorescence bracts (floret lemmas and spikelet glumes) are not suppressed in the grasses, all bracts subtending inflorescence branches and spikelets are suppressed. This selective bract suppression in grasses is morphologically distinct from other angiosperms lineages such as the Brassicaceae, where bracts subtending both flowers and inflorescence branches are generally suppressed (*4*). Positionally specific suppression of bracts is a morphological innovation of the grass family as close outgroups in Poales do not suppress bracts at any position in their inflorescence (*3*).

Analysis of several bract suppression mutants in maize has provided key insights into the molecular regulation of bract suppression in the grass family. These mutants include the *tassel sheath 1-5* (*tsh1-5*) loci (*3*), of which two have been cloned. *tsh1* encodes a GATA domain zinc-finger transcription factor (*3*), while *tsh4* encodes a *SQUAMOSA PROMOTER BINDING PROTEIN* (*SBP*) transcription factor (*5*). The dominant *Cg1* also displays de-repressed bracts and encodes a microRNA that targets *tsh4* and related *SBP* family members (*6*). Two *SBP* genes, *unbranched2* and *3*, are closely related to and are redundant with *tsh4* to regulate bract suppression and inflorescence branching (*7*). Other *tsh* loci have not yet been cloned, but it is intriguing that genes from at least two unrelated transcription factor families are required for bract suppression in maize, suggesting that a complex transcriptional network for bract suppression evolved in the grass family.

Bract suppression in eudicots and grasses is likely controlled by distinct genes. In *Arabidopsis thaliana* (arabidopsis), bract suppressing genes include *LEAFY (8)* and *BLADE ON PETIOLE* (*BOP*) *1* and *BOP2* (*9, 10*). Of these, *LFY* plays a major role, which appears to be conserved in eudicot lineages that independently evolved bract suppression, including Solanaceae and Fabaceae (*11*–*13*). However, loss-of-function mutants for orthologs of *LFY* and other eudicot bract suppression genes show no bract growth defects in the grasses (*14*–*16*). Similarly, grass bract suppression genes have no bract suppression role in arabidopsis. While *tsh4* and orthologous *SBP* genes have a conserved phase transition function in both eudicots (*17*) and monocots (*5, 7*), *SBP-like* (*SPL*) *9* and *SPL15* genes, the arabidopsis orthologs of *tsh4*/*ub2*/*ub3*, do not influence bract suppression (*18*). A comparison of *tsh1* function across eudicots and grasses reveals even more divergence in their bract suppression pathways. Knockouts of the arabidopsis *tsh1* orthologs *HANABA TARANU* (*HAN*), *HAN-like1* and *HAN-like2* show defects in floral organ initiation and separation, and embryo patterning consistent with a boundary domain function (*19*–*21*). These boundary phenotypes are not evident in *tsh1* mutants, indicating a dramatic shift in *HAN* vs. *tsh1* function at some point in angiosperm evolution. Why grasses evolved a novel and complex bract suppression network that is morphologically targeted to only a subset of inflorescence branching events is not clear.

While the developmental and evolutionary role of bract suppression is still an open question, a proposed explanation is that the emerging bract competes with the adjacent meristem for cells and growth factors. Bract suppression, in this interpretation, diverts limited growth resources away from leaves and toward meristem growth and branching after the floral transition (*22*–*25*). In support of this hypothesis, de-repressed bract growth is correlated with reduced branching in maize *tsh1* and *tsh4* mutants (*3, 5*). This correlation is not complete, however, as *tsh* mutants will often form bracts without affecting the determinacy of their adjacent meristem (*3*). Conversely, *ub2* and *ub3* affect branching but not bract growth, despite their localization to the bract primordium and redundant function when combined with *tsh4* to suppress bracts (*7*). The partial decoupling of bract growth from branch suppression raises the possibility the suppressed bract and the meristem it subtends interact in a manner beyond mere competition for resources. One possibility is that *tsh1* and *tsh4* genes are involved in regulation of grass branch architecture as part of a suppressed bract signaling center that non-autonomously regulates the determinacy of the branch meristem (*26*).

Considering the importance of meristem determinacy to inflorescence architecture and yield traits in domesticated cereals, we sought to better understand the contribution of bract suppression to inflorescence development and the bract transcriptional networks regulated by *tsh1* and *tsh4*. Here, we report our investigation of genetic interactions and transcriptional changes associated with these bract suppression mutants. We identify a core transcriptional network involving *tsh1, tsh4* and the boundary domain gene *liguleless2* (*lg2*), that jointly regulate both bract suppression and inflorescence branch meristem determinacy. We detect a series of duplications in the grass *tsh1* gene lineage that preceded the recruitment of this boundary domain gene to the suppressed bract. Our results suggest that the phase change regulator *tsh4* recruited *tsh1* and *lg2* to a targeted role in inflorescence development, and that bract suppression indirectly resulted from recruiting these boundary domain genes to form a novel signaling center that promotes branch meristem indeterminacy in grasses.

## Results

### *tsh4* acts synergistically with *tsh1* to regulate bract suppression and branch meristem indeterminacy

In an ongoing effort to characterize additional *tsh* loci, we identified a novel allele of *tsh4* (described previously as *tsh2*) containing a *Mutator* transposon insertion in the first intron, which we designated *tsh4-rm* (fig. S1, A and C). In a separate screen for genetic modifiers of the weak *tsh1-2* allele, we isolated a semidominant *tsh1* enhancer that was also allelic to *tsh4* (*tsh4-ent*355*; fig. S1, B and C), which segregated as a single, recessive locus in the absence of *tsh1-2*. In addition, we identified a large deletion paired with a *Mu* transposon insertion as the causative lesion in the *tsh1-ref* allele (fig. S1D).

The dramatic enhancement of the weak *tsh1-2* phenotype by *tsh4-ent*355* suggests a synergistic interaction. To more fully investigate the nature of this interaction, we measured both tassel branching and bract suppression in a F2 population segregating *tsh4-rm* and *tsh1-ref*, each introgressed ≥ 5X to the reference B73 genetic background. Introgression of *tsh4-rm* into B73 significantly suppressed the phenotype (compare fig.S1A with Fig.4D) indicating that natural modifiers in B73 ameliorate the phenotypic severity of *tsh4* mutants. Nevertheless, *tsh4* and *tsh1* showed a consistent and striking synergistic interaction. As shown in Fig.1 and Table 1, *tsh1 tsh4* double mutant tassels produce no long tassel branches and compared to *tsh1* and *tsh4* single mutants, *tsh1 tsh4* double mutant tassels also have a non-additive increase in the percentage of solitary spikelets, empty nodes, and nodes with subtending de-repressed bracts. The phenotypic enhancement was not limited to the double mutant as *tsh1*/*+*; *tsh4*/*tsh4* and *tsh1*/*tsh1*; *tsh4*/*+* individuals had fewer branches and more bract growth than *tsh1* or *tsh4* single mutants.

**Fig. 1.**
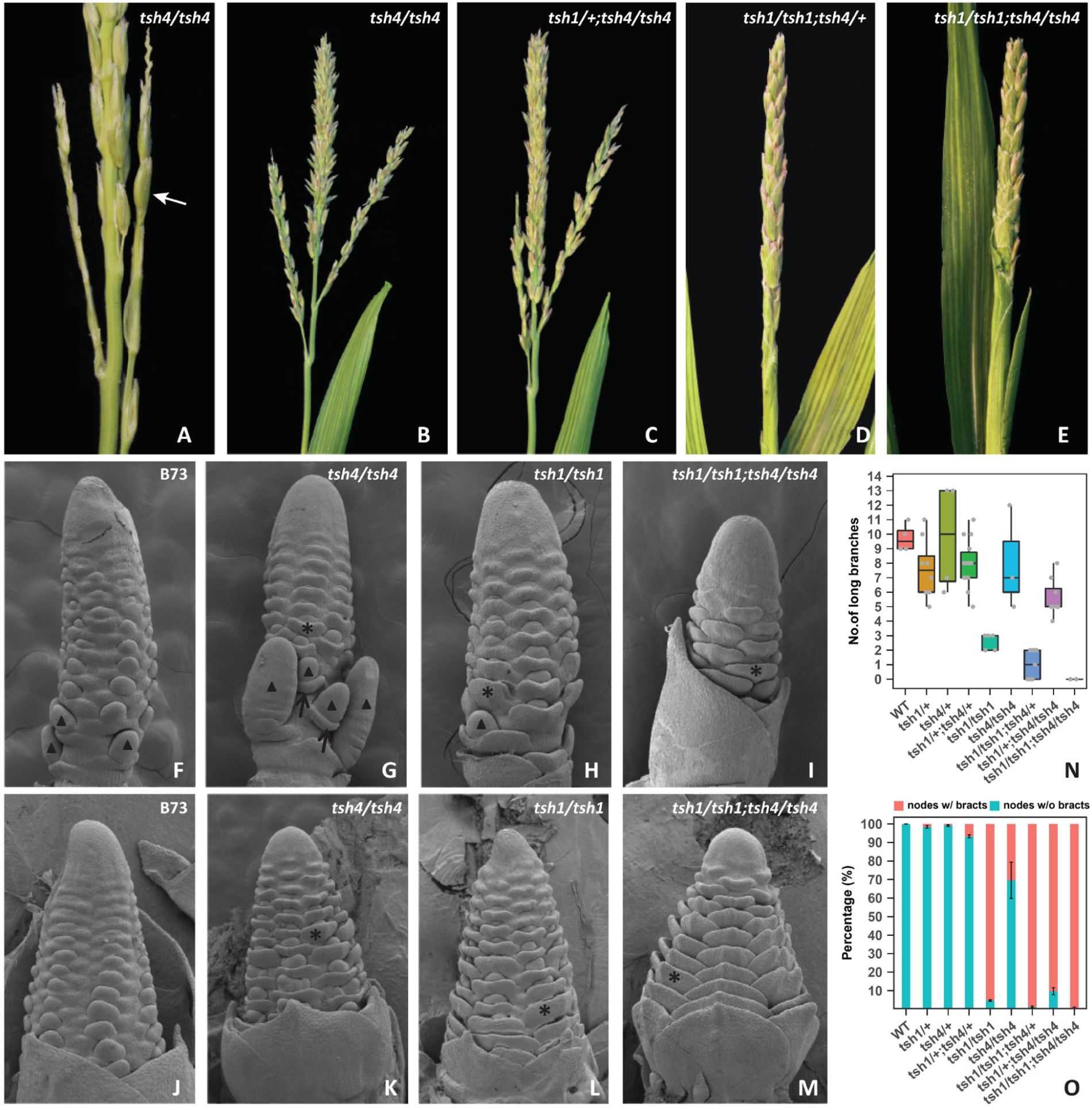
*tsh1* and *tsh4* act synergistically to regulate bract suppression and inflorescence branching. (**A** to **E)** Tassel phenotype of plants in *tsh1-ref/+*: *tsh4-rm/*+ (B73) segregating population showing progressive enhancement of *tsh4* (A) as *tsh1* function is progressively removed. **(F** to **M**) Scanning electron microscopy of tassel (F-I) and ear (J-M) inflorescence primordia of wildtype B73 (F, J), *tsh4* (G, K), *tsh1* (H, L), and *tsh1 tsh4* double mutant (I, M). (**N** to **O**) Quantification of branching (N) and bract (O) phenotype in all segregating genotypes. Arrow in A designates a bract, as do asterisks (*) in F to M. Arrowheads in F to M indicate long tassel branches.

**Table 1.** Phenotypic characterization of *tsh1*/*+*; *tsh4*/+ segregating population. A *tsh1-ref*/+, *tsh4-rm*/+ segregating population containing 57 plants was grown in Spanish fork, UT in 2018 and PCR genotyped. Mature tassels from all genotyped plants were inspected manually for branch and bract growth. In this table we report the mean value for each genotype with the s.e.m. in parentheses. Statistical significance was determined by one-way ANOVA (significance level = 0.05) followed by post-hoc comparisons to determine if statistically significant differences exist between the *tsh1* single mutant (*tsh1*/*tsh1* homozygote) and the *tsh1 tsh4* mutants (*tsh1*/*tsh1*; *tsh4*/+ mutant or *tsh1*/*tsh1*; *tsh4*/*tsh4* mutant), or between *tsh4* single mutant (*tsh*4/*tsh4* homozygote) and the *tsh1 tsh4* mutants (*tsh1*/+; *tsh4*/*tsh4* mutant or *tsh1*/*tsh1*; *tsh4*/*tsh4* mutant) by using Fisher’s LSD method. significant difference, **P < 0.01, ***P < 0.001, ns, not significant.

| Genotype | No. of long branches | % nodes with long branches | % nodes with paired spikelets | % nodes with solitary spikelets | % empty nodes | % nodes with subtending bracts |
| --- | --- | --- | --- | --- | --- | --- |
| WT | 9.750(0.479) | 0.040(0.005) | 0.907(0.024) | 0.053(0.019) | 0.000(0.000) | 0.000(0.000) |
| *tsh1/+* | 7.625(0.730) | 0.030(0.002) | 0.941(0.012) | 0.029(0.010) | 0.000(0.000) | 0.015(0.008) |
| *tsh4/+* | 9.750(1.887) | 0.039(0.004) | 0.904(0.008) | 0.057(0.007) | 0.000(0.000) | 0.008(0.005) |
| *tsh1/+;tsh4/+* | 8.000(0.432) | 0.032(0.002) | 0.907(0.008) | 0.061(0.006) | 0.001(0.000) | 0.067(0.008) |
| *tsh4/tsh4* | 8.000(2.082) | 0.030(0.005) | 0.792(0.055) | 0.170(0.050) | 0.009(0.003) | 0.304(0.098) |
| *tsh1/+;tsh4/tsh4* | 5.625(0.460)<sup>ns</sup> | 0.032(0.003)<sup>ns</sup> | 0.463(0.037)<sup>***</sup> | 0.498(0.038)<sup>***</sup> | 0.007(0.002)<sup>ns</sup> | 0.903(0.019)<sup>***</sup> |
| *tsh1/tsh1* | 2.600(0.245) | 0.021(0.002) | 0.753(0.025) | 0.216(0.024) | 0.010(0.003) | 0.952(0.004) |
| *tsh1/tsh1 tsh4/+* | 1.000(0.333)<sup>ns</sup> | 0.010(0.003)<sup>**</sup> | 0.591(0.033)<sup>***</sup> | 0.360(0.032)<sup>***</sup> | 0.040(0.007)<sup>***</sup> | 0.987(0.004)<sup>ns</sup> |
| *tsh1/tsh1;tsh4/tsh4* | 0.000(0.000)<sup>***</sup> | 0.000(0.000)<sup>**</sup> | 0.289(0.036)<sup>***</sup> | 0.543(0.043)<sup>***</sup> | 0.168(0.007)<sup>***</sup> | 0.994(0.006)<sup>***</sup> |

In order to investigate the early ontogeny of the branching and bract growth defects, we examined tassel and ear primordia of *tsh1* and *tsh4* single and double mutants by scanning electron microscopy (Fig. 1, F-M). While most nodes were associated with a de-repressed bract in *tsh1*, bract growth was more pronounced in the double mutant, particularly in the ear. Interestingly, while derepressed bracts clearly subtend axillary meristems in *tsh1* and *tsh4* single mutants, *tsh1 tsh4* double mutants lack any obvious axillary meristems at these early stages (Fig. 1M). These results confirm that *tsh1* and *tsh4* act redundantly to suppress bract growth and promote meristem initiation.

### *tsh1* and *tsh4* regulate diverse pathways involved in meristem and leaf development, hormone signaling, and boundary domains

Transcripts of both *tsh1* and *tsh4* are localized to the bract primordium from the earliest stages of bract initiation (*3, 5, 6*). However, the molecular processes regulated by these genes within this very narrow domain is unclear. To identify transcriptional changes associated with *tsh1-* and *tsh4*-mediated bract suppression, we generated RNA-seq transcriptomes of laser-microdissected (LM) bract primordia of the wildtype (B73), *tsh1, tsh4* and *tsh1 tsh4* double mutants (DM). Specifically, we collected cells from ear bract primordia where the bract ridge, but not the adjacent meristem is visible (Fig. 2A). Principal Component Analyses (PCA) confirmed that the biological replicates were highly correlated within genotypes (fig. S2). In addition, *tsh1* and *tsh4* transcript levels were significantly down-regulated in their respective mutants (fig. S3). To confirm the tissue specificity of our LM, we investigated the known suppressed bract marker *Zea yabby14* (*Zyb14*) *(3)*, as well as the meristem-specific *Knotted1* (*Kn1*), which is strongly down-regulated in the suppressed bract (*27*). As expected, there was significant enrichment of *Zyb14* and reduction of *Kn1* compared to their expression in LM shoot apical meristem tissue (fig. S4). Thus, our transcriptomes are reliable and will likely uncover transcriptional changes associated with bract suppression downstream of *tsh1* and *tsh4*.

**Fig. 2.**
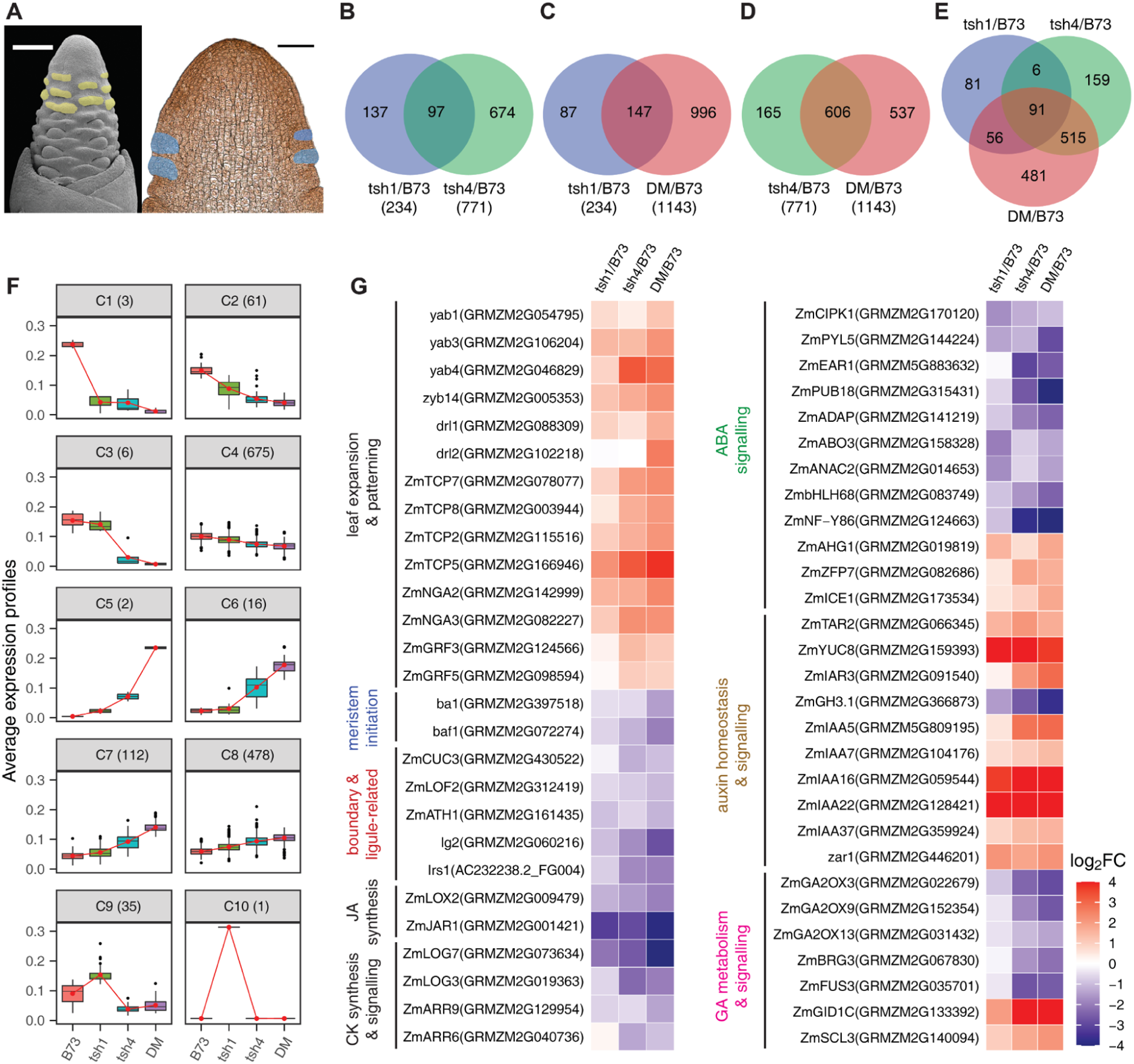
Transcript profiling of laser-microdissected suppressed and growing ear bract primordia. (**A**) SEM (left) and thin section (right) of young ear primordium indicating the cells targeted for laser capture (yellow and blue respectively) scale bars = 200 µm (left) or 100 µm (right). (**B** to **E**) Venn diagram of common and unique differentially regulated bract genes in *tsh1, tsh4* single and *tsh1 tsh4* double mutants (DM). (**F**) Ten co-expression clusters (C1–C10) were identified from the 1389 genes that were differentially expressed between B73 and the *tsh* mutants. Connected red lines correspond to the mean expression profiles for each cluster. The vertical bars define the upper or lower quartile, and dots outside the bars indicate outliers. (**G**) Genes with well-documented function in leaf expansion and patterning, meristem initiation, boundary and ligule establishment, and hormone metabolism/signaling were differentially expressed in *tsh* mutants compared to that in the wildtype.

In total, 31.8% (20,168) of maize annotated genes (AGPv3_5b+) were expressed in at least one sample (table S1). Of these, 6.9% (1,389) were differentially expressed genes (DEGs) between B73 and at least one of the mutants (Fig. 2B-2E, table S2-S5). Over three times more genes were differentially expressed (DE) in *tsh4* (771) compared to *tsh1* (234), suggesting that *tsh1* regulates a narrower set of downstream genes (Fig. 2B). DEGs in the *tsh1 tsh4* double mutant included the majority of the genes that were differentially expressed in *tsh1* and *tsh4* individually (60% and 78%, respectively) but also included an additional 481 DEGs not present in either single mutant, consistent with the synergistic phenotype of *tsh1 tsh4* (Fig. 2C-2E). The PCA was largely consistent with these conclusions as PC1 (63% of variance) is mostly explained by *tsh4* while PC2 (17% of variance) is explained by *tsh1*, indicating partially non-overlapping roles for *tsh1* and *tsh4*, but with more dramatic impact from *tsh4* (fig. S2). While *tsh1* and *tsh4* have distinct DEG profiles, a significantly larger portion of *tsh1* DEGs were shared with *tsh4* than vice versa (41% vs 12%) (Fig. 2B). This raises the possibility that *tsh1* functions downstream of *tsh4*. Therefore, we examined expression of *tsh1* in *tsh4* single mutants and vice versa. While *tsh4* expression in *tsh1* single mutants is similar to B73 controls, *tsh1* is significantly decreased in *tsh4* mutants (fig. S3), consistent with *tsh4* functioning upstream of *tsh1*.

To gain further insight into the molecular processes associated with bract suppression, we employed a combination of K-means clustering and Gene Ontology (GO) classification enrichment analysis on DEGs in at least one *tsh* mutant. K-means analysis identified ten unique clusters of co-expressed genes across the different genotypes (Fig. 2F, table S6a and S6b). Clusters 1-4 showed similar patterns of decreased expression in the mutants; genes in these clusters likely function downstream of *tsh1* and *tsh4* to repress bract growth. Clusters 5-8 showed the opposite trend, with increased expression in the mutants; genes in these clusters likely promote bract outgrowth. A common trend observed in clusters 1-8 was a more dramatic response in *tsh4* mutants compared to *tsh1*, with the strongest response in the double mutant, consistent with *tsh4* acting upstream of *tsh1*. In contrast, clusters 9 and 10 include a small set of genes that were upregulated only in *tsh1*. We performed GO enrichment analysis for genes in clusters 1-4 and 5-8 (fig. S5). Overall, these clusters are enriched for genes involved in the regulation of gene expression, organ developmental processes (leaf, root, flower, inflorescence, meristem), cell growth and differentiation, and hormone metabolism and signaling. Interestingly, some enriched GO categories were exclusive to the upward vs. downward trending expression in clusters 1-4 vs. 5-8. In particular, genes involved in inflorescence development, gibberellin metabolic process, response to abscisic acid, response to ethylene, extracellular matrix assembly and dormancy were unique to clusters 1-4, while genes involved in leaf morphogenesis, auxin metabolic process, auxin transport and auxin-activated signaling pathways were only enriched in clusters 5-8. Clusters 9-10 contain only 36 genes with no significantly enriched GO terms.

A closer look at individual DEGs revealed several genes with known roles in leaf expansion and patterning as well as branch meristem initiation and determinacy. These results were largely expected based on the bract growth and branch suppression phenotypes of *tsh1* and *tsh4*. In addition to these genes, we were particularly interested in genes involved in hormone metabolism/signaling and boundary formation given the known function of *HANABA TARANU* (*HAN*, the arabidopsis *tsh1* ortholog) as a boundary gene that regulates hormone dynamics (*19*–*21*).

#### Leaf expansion and patterning

Multiple transcription factor families with well-documented roles in leaf development and patterning were differentially regulated in *tsh* mutants, consistent with the bract outgrowth phenotype of these mutants (Fig. 2G). This included upregulation of multiple *YABBY* transcription factors (*yab1, yab3, zyb14, drl1* and *drl2*), which have critical roles in both leaf expansion and polarity (*28, 29*). Additionally, *TEOSINTE BRANCHED1, CYCLOIDEA, PCF1* (*TCP*) transcription factors including both class I (*ZmTCP7* and *8*) and class II TCP-CIN (*ZmTCP2* and *5*) orthologs were upregulated in *tsh*. While the function of class I TCPs are poorly understood, they are thought to regulate cell division (*30*). Class II-TCP orthologs, however, have a well-documented role in leaf development (*31*), and interact with *NGATHA* (*NGA*) genes to promote cell differentiation and leaf expansion (*32*). Interestingly *NGA2* and *3* are also upregulated in *tsh* bracts. Finally, two *GROWTH-REGULATING FACTOR* transcription factors (*GRF3* and *5*) are upregulated. These genes regulate cell proliferation in Arabidopsis (*33*) and are enriched in the actively dividing regions of the maize leaf (*34*), consistent with a role in early leaf expansion.

#### Meristem initiation

Inflorescence branching is controlled by genes that regulate axillary meristem initiation and determinacy. In maize, the transcription factors *barren stalk1* (*ba1*) and *barren stalk fastigiate1* (*baf1*) are required for axillary meristem initiation in the inflorescence (*35, 36*). Both genes are significantly downregulated in *tsh* mutants (Fig. 2G), consistent with the reduced and delayed branch meristem initiation in *tsh* mutants. *ba1* and *baf1* are not expressed in the suppressed bract, but in an adjacent boundary domain adaxial to the axillary meristem. It is possible that adjacent cells expressing *ba1/baf1* were sampled inadvertently from the margins of captured bract cells during laser microdissection.

#### Hormone homeostasis and signaling

Lateral organ growth is regulated by a complex interplay of hormone signaling. In light of this it is not surprising to find a number of hormone signaling genes that were differentially regulated. Specifically, our data consistently indicate that Abscisic Acid (ABA), auxin, and Gibberellic Acid (GA) signaling were altered.

ABA signaling components including maize orthologs of *CBL-INTERACTING PROTEIN KINASE 1* (*CIPK1*), *PYRABACTIN RESISTANCE 1-LIKE 5* (*PYL5*), *ENHANCER OF ABA CO-RECEPTOR 2* (*EAR1*), *PLANT U-BOX 18* (*PUB18*), *ARIA-INTERACTING DOUBLE AP2 DOMAIN PROTEIN* (*ADAP*), *ABA OVERLY SENSITIVE MUTANT 3* (*ABO3*), *ARABIDOPSIS NAC DOMAIN CONTAINING PROTEIN 2* (*ANAC2*), *BASIC HELIX-LOOP-HELIX DNA-BINDING FAMILY PROTEIN* (*bHLH68*) and *NUCLEAR FACTOR Y, SUBUNIT B6* (*NF-YB6*) were down-regulated in *tsh* mutants (Fig. 2G), suggesting that *tsh1* and *tsh4* promote ABA signaling in the suppressed bract. Consistent with this, the orthologs of three negative regulators of ABA signaling, including *ABA-HYPERSENSITIVE GERMINATION 1* (*AHG1*), *ZINC FINGER PROTEIN 7* (*ZFP7*) (*37*) and *INDUCER OF CBF EXPRESSION 1* (*ICE1*) *(38)*, were up-regulated in *tsh* mutants (Fig. 2G). As ABA is associated with dormancy and growth inhibition in multiple developmental contexts(*39*), *tsh1/tsh4* may promote ABA signaling to inhibit bract outgrowth.

Auxin is another crucial regulator of lateral organ initiation and outgrowth (*40*). The auxin biosynthesis genes *TRYPTOPHAN AMINOTRANSFERASE RELATED 2* (*TAR2*) and *YUCCA 8* (*YUC8*) and an auxin conjugate hydrolase *IAA-ALANINE RESISTANT 3* (*IAR3*) were upregulated in *tsh* mutants, whereas an auxin inactivation gene IAA-amido synthetase *GH3*.*1* was downregulated compared to wildtype (Fig. 2G). This suggests that *tsh1* and *tsh4* may inhibit auxin production in the suppressed bract primordium. Consistent with this, auxin-responsive Aux/IAA family transcription factors including *IAA5, IAA7, IAA16, IAA22* and *IAA37* were upregulated in *tsh* mutants. Similarly, the auxin-inducible gene *zar1*, a positive regulator of cell proliferation and lateral organ size (*41*), was upregulated, consistent with the bract outgrowth phenotype in *tsh* mutants.

GA promotes organ growth by inducing cell division and elongation (*42*). We found that the GA inactivation enzymes *ZmGA2OX3, ZmGA2OX9* and *ZmGA2OX13* and GA response inhibitors *BOI-RELATED GENE 3* (*BRG3*) (*43*) and *FUSCA3* (*FUS3*) (*44*) were downregulated in *tsh* mutants (Fig. 2G). Conversely, orthologs of a GA receptor *GA INSENSITIVE DWARF1C* (*GID1C*) and the positive regulator of GA signaling *SCARECROW-LIKE 3* (*SCL3*) (*45*) were upregulated. Taken together, these data indicate that *tsh1* and *tsh4* potentially inhibit GA signaling to suppress bract growth.

In addition to ABA, auxin and GA, several genes involved in the metabolism and/or signaling of Jasmonic Acid (JA) and cytokinin (CK) were differentially expressed between wildtype and *tsh* bracts (Fig. 2G). These included orthologs of the JA biosynthesis genes *LIPOXYGENASE 2* (*LOX2*) and *JASMONATE RESISTANT 1* (*JAR1*), which were downregulated in *tsh* mutants. Cytokinin-activating enzymes (*LONELY GUY 3* (*LOG3*) and *LOG7*) and CK signaling components (*TYPE-A RESPONSE REGULATOR 6* (*ARR6*) and *ARR9*) were downregulated in *tsh* mutants suggesting attenuated CK biosynthesis and signaling in the *tsh* bracts. Taken together, our transcriptomic analysis suggests that *tsh1* and *tsh4* promote ABA, JA and CK signals while attenuating auxin and GA signals in the bract primordium.

#### Boundary Domain and Ligule-associated genes

Boundary domain genes were first described in eudicots where they separate and promote morphogenesis of determinate lateral organs from indeterminate meristems (*46*). Less is known about boundary formation in the monocots, and grasses in particular appear to have a novel boundary in the leaf known as the ligule that separates proximal sheath from distal blade compartments (*47*). Interestingly, mutants lacking a ligule also have defects in tassel branching (*48, 49*) similar to *tsh* mutants. Furthermore, the rice *tsh1* ortholog *NECKLEAF1* is expressed in the ligule (*50*), and *tsh1* expression was shown to be enriched in the preligular band compared to adjacent blade and sheath tissue (*51*). Considering these correlations between boundary genes, ligules, and tassel branching, we were curious if boundary or ligule genes were differentially regulated in *tsh* mutants.

We found several genes with documented boundary domain functions in arabidopsis including orthologs of *CUP SHAPED COTYLEDON3* (*CUC3*) (*52*), *LATERAL ORGAN FUSION2* (*LOF2*) (*53*) and *ARABIDOPSIS THALIANA HOMEOBOX GENE1* (*ATH1*) (*54*), all of which were downregulated in the *tsh* mutants (Fig. 2G).

In addition to canonical boundary domain genes, we found that *lg2*, and *liguleless related sequence1* (*lrs1*), a paralog of *lg2*, were downregulated in *tsh* mutants (Fig. 2G). Given the known connections between bracts and ligules, we were curious if transcriptional changes associated with ligule development were similarly present in our dataset. In order to investigate this, we compared our bract transcriptome with a published LM expression profile of cells early in ligule specification (*51*). Compared to neighboring pre-blade and pre-sheath cells, the preligule was enriched for 619 genes (*51*). Among these 619 DEGs, a significant portion (141 genes, 22.8%, hypergeometric test p-value = 3.209e-99) were also differentially expressed between wildtype and *tsh* bract primordia (fig. S6 and table S7). The same study also identified 96 DEGs between wild type and *liguleless1* (*lg1*), which is another gene required for ligule development (*55*). Interestingly, a significant proportion of these (24 genes, 25%, hypergeometric test p-value = 1.967e-15) were also differentially expressed in *tsh* mutants (fig. S6). In addition, 21 (87.5%) out of these 24 shared DEGs exhibited a similar trend of expression change in *lg1* and *tsh* mutants (table S8). Taken together, these results confirm that many transcriptomic changes associated with ligule determination are also associated with bract suppression.

### TSH4 binds regulatory DNA of *tsh1* and *lg2*

The downregulation of *tsh1* in *tsh4* mutants suggests that *tsh4* is upstream of *tsh1* in a bract suppression network. In order to test if this interaction is direct, we used a previously developed TSH4 antibody (*5*) to perform Chromatin Immuno Precipitation (ChIP). The TSH4 antibody was used to precipitate chromatin from ≤5mm ear primordia, and enrichment was quantitated by qPCR amplification of five regions across the *tsh1* gene (Fig. 3A). Compared with chromatin purified by IgG negative control, strong enrichment was observed for two regions (b and c) in the *tsh1* promoter approximately 1kb upstream of the transcription start site. In addition, we found these two regions were also bound by TSH4 in a TSH4 DAP-seq dataset (*56*) demonstrating that *tsh1* is a direct binding target of TSH4 in vivo. While *tsh1* levels are strongly reduced in *tsh4* mutants, some residual expression suggests that additional factors are required to initiate *tsh1* in the suppressed bract consistent with the significantly milder bract suppression phenotype of *tsh4* compared to *tsh1*.

**Fig. 3.**
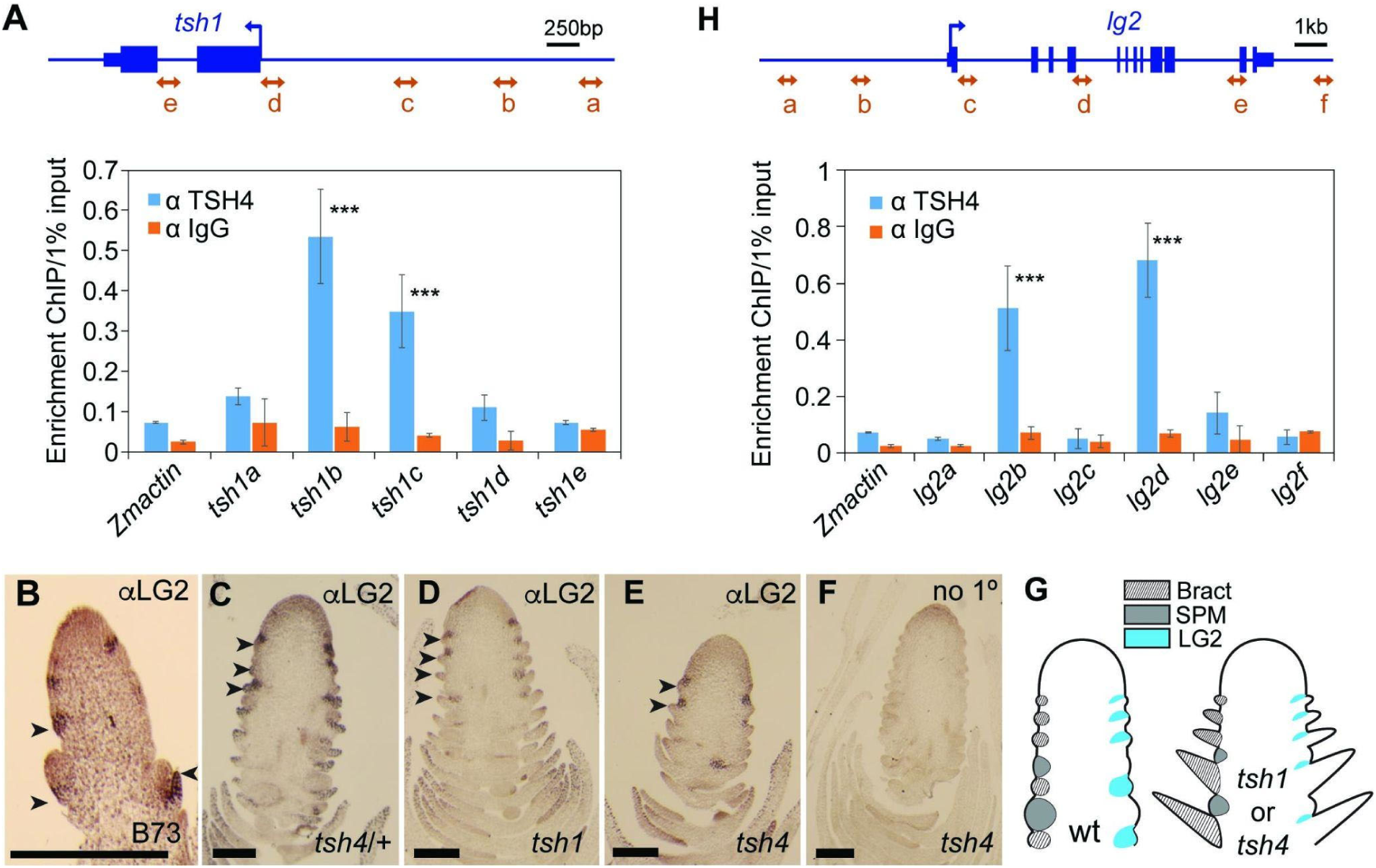
TSH4 binds promoters of *tsh1* and *lg2* and is necessary for suppressed bract expression of *lg2*. (**A**) anti-TSH4 ChIP using primers designed to five locations (arrows a-e) in the *tsh1* genomic region. Bar graph shows enrichment of an anti-TSH4 ChIP compared to an anti-IgG control. Significant enrichment was found for promoter regions *tsh1b* and *tsh1c*. (**B** to **E**) Immunolocalization of LG2 on a wildtype (B73) tassel primordium (B), and ear primordia of *tsh4/+* (C), *tsh1* (D), and *tsh4* (E). (**F**) Immunolocalization control without primary anti-LG2 antibody shows that the bract and boundary domain localization in B to E is specific to anti-LG2. (**G**) Summary of LG2 localization in wt (left) and *tsh1* or *tsh4* mutants (right). SPM=spikelet pair meristem. (**H**) anti-TSH4 ChIP as in (A), using primers designed to six regions (a-f) of the *lg2* genomic region. Significant enrichment (*** indicates P < 0.001 by Student’s t test.) was observed for the promoter (*lg2b*) and the fourth intron (*lg2d*). Arrowheads in B to E indicate bract primordia.

Among the genes differentially regulated by *tsh1* and *tsh4*, is *lg2*, a transcription factor required for ligule development(*57*). Considering the reduced branching of *lg2* mutants (*48*), *lg2* may interact with *tsh1* and *tsh4* in their branch promotion roles. While *lg2* mutants were not originally described as having any bract suppression defects, we noticed that both tassels and ears of *lg2* mutants stochastically produce large bracts with low penetrance (fig. 7A and B), further pointing to a connection of bract regulation and branch meristem determinacy.

To investigate the localization of LG2 during tassel development, we raised an antibody specific to LG2 (fig. 7C), and used this for immunolocalization at early stages of inflorescence development. We found that LG2 localizes to a broad domain that includes both the suppressed bract and the boundary between the suppressed bract and the adjacent meristem (Fig. 3B). The localization of LG2 overlaps with the bract-expression of *tsh1* and *tsh4* (*3, 5*). Given the reduced *lg2* mRNA levels in both *tsh1* and *tsh4*, we hypothesized that these regulators of bract suppression are necessary to promote LG2 protein accumulation within the suppressed bract. Indeed, LG2 expression is reduced or absent from the suppressed bract region of *tsh1* and *tsh4* mutants, while largely maintained in the narrow boundary between the suppressed bract and the adjacent meristem (Fig. 3B-3G), indicating that *lg2* is downstream of *tsh1* and *tsh4* within the suppressed bract.

We asked if transcriptional regulation of *lg2* could be explained by direct binding of TSH4 to its promoter. Through ChIP qPCR across the *lg2* genic region, we identified two strong binding regions located in the proximal promoter and the fourth intron respectively (Fig. 3H). Of these two apparent binding sites, the fourth intron was also bound by TSH4 in a TSH4 DAP-seq dataset (*56*). These results suggest that *lg2* is a transcriptionally modulated direct target of TSH4. Altogether, our results reveal that *tsh4* is a positive regulator of *tsh1* and *lg2*. In addition, *tsh1* appears to positively regulate *lg2* independent of and redundantly with *tsh4*.

### *lg2* interacts synergistically with *tsh1* and *tsh4* to regulate bract suppression, and branch meristem determinacy

Our observation that *tsh1* and *tsh4* redundantly promote *lg2* in the suppressed bract, raises the possibility that these factors cooperate in bract suppression and/or inflorescence branch meristem determinacy. To assess any genetic interaction of *lg2* with *tsh1* or *tsh4*, we generated *tsh1*/*+*; *lg2*/*+* and *tsh4*/*+*; *lg2*/*+* segregating populations and compared the tassel phenotype of each individual genotype in the populations (Fig. 4 and table S9-S10). Removing a copy of *lg2* (*lg2*/+) from a *tsh1* or a *tsh4* homozygous background significantly reduced long basal branches (Fig. 4B and E). Similarly, removing a copy of *tsh1* (*tsh1*/+) or *tsh4* (*tsh4*/+) from a homozygous *lg2* background reduced long branches. Double mutants of *tsh1 lg2* and *tsh4 lg2* completely lacked long branches. Similar shifts of paired to solitary spikelets or nodes lacking any spikelet were observed for each of these genotypes (table S9 and S10). While *lg2* only rarely produces tassel bracts, removing a copy of *tsh1* (*tsh1/+*) or *tsh4* (*tsh4/+*) from a *lg2* homozygous background resulted in consistent bract production, and the double mutants (both *tsh1 lg2* and *tsh4 lg2*) had a significant increase in tassel nodes with bracts (Fig. 4C and F). These synergistic interactions are consistent with redundant and co-operative roles for *lg2* with both *tsh1* and *tsh4* in suppressing inflorescence bract growth and promoting branch meristem indeterminacy. We also noticed that while bracts often subtended branches in *tsh* mutants, reduced branches (solitary spikelets and empty nodes) were not always subtended by bracts (fig. S8 and table S11-S12), consistent with a role for *tsh1* and *tsh4* in promoting branch indeterminacy independent from their role in bract suppression. These genetic interactions further confirm that *lg2* functions in a common regulatory network with *tsh1* and *tsh4*, and underscore the importance of boundary domain genes in bract suppression and associated branch meristem determinacy.

**Fig. 4.**
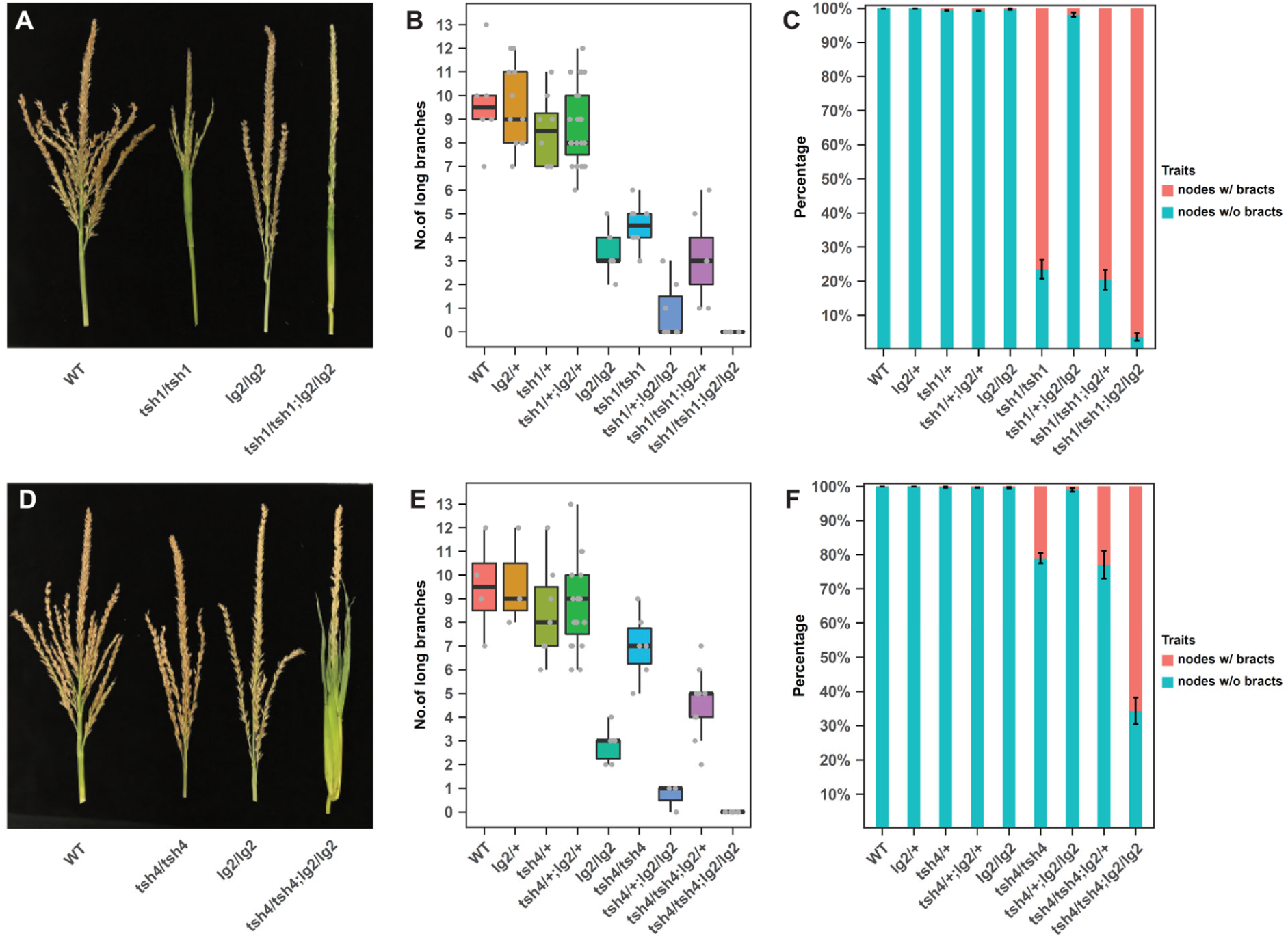
Synergistic *tsh1 lg2* and *tsh4 lg2* interactions promote bract growth and branch repression. (**A** to **C**) *tsh1 lg2* and (**D** to **F**) *tsh4 lg2* genetic interactions. Mature tassel phenotypes (A, D) show an enhancement of bract growth and reduced tassel branching in double mutants, which is further confirmed in a quantification of the number of long branches (B, E) and nodes (i.e., branching sites of the inflorescence) with bracts (C, F).

### *tsh1* was likely recruited from an ancestral boundary domain function

The expression pattern and mutant phenotypes of *tsh1* and orthologous genes in the grasses (*3, 58*) sharply contrast with arabidopsis *HAN* (*59*). One possible explanation for this divergence is that the boundary domain expression and function in arabidopsis is ancestral, and that the grass *NECKLEAF1, tsh1, THIRD OUTER GLUME* (*NTT*) clade evolved a novel expression and function related to bract suppression and branch promotion. Given the known duplications of the eudicot *HAN-like* and grass *NTT* genes (*3, 19*), such neofunctionalization is a possibility. As a first step toward investigating the functional divergence of the *HAN-NTT* gene family, we reconstructed their phylogeny focusing in particular on the history of duplications in the grasses (Poaceae) and broader Poales (Fig. 5A). Our analysis revealed that the *HAN-NTT* subfamily of GATA domain transcription factors (i.e., those containing both a GATA and HAN domain) are present throughout the land plants, including the liverwort *Marchantia polymorpha* and the moss *Physcomitrium patens*. Within the eudicots, multiple duplications are apparent and, although our sampling was not sufficient to resolve the timing for each of these, none showed evidence of dating to deep nodes.

**Fig 5.**
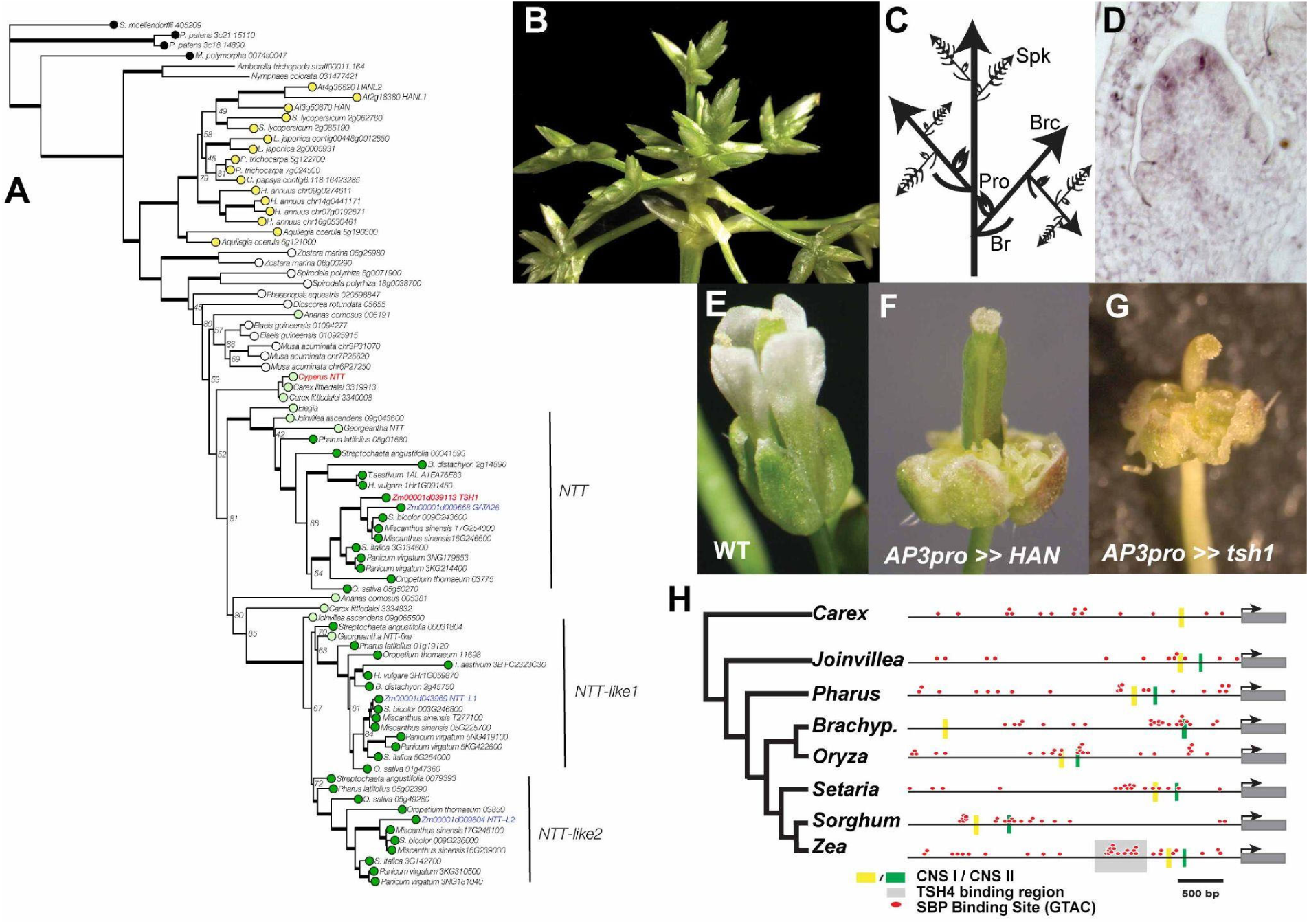
The NTT lineage in the grass family may result from neofunctionalization and maintains an ancestral boundary domain activity. (**A**) Phylogeny of the *HAN-NTT* gene family. (**B** to **C**) *Cyperus* inflorescence shows prominent bracts and prophylls associated with all inflorescence branching events. (**D**) In situ localization of *Cyperus NTT* in inflorescence shows *Cyperus NTT* expresses in the boundary domain separating the spikelet meristem from lateral florets. (**E** to **G**) Phenotypes of arabidopsis florets with ectopic expression of *HAN* (F) or *Tsh1* (G) under *AP3* promoter. WT, wildtype control (E). (**H**) Distribution of putative SBP binding sites in 5’ promoters of *NTT* genes from grasses and close outgroups. CNS = Conserved Non-coding Site.

In the monocots, we identified a well-supported clade of Poales *NTT* genes. Within this Poales clade we identified a duplication resulting in two distinct clades: *NTT* and *NTT-like1/2*. Members of the *NTT-like* clade are found in all Poales lineages sampled including *Joinvillea* and *Carex* (Cyperaceae) and *Ananas* (Bromeliaceae). However, the *NTT* clade includes no representative from *Ananas* or Cyperaceae, which are sister to the larger *NTT/NTT-like* clade, although the placement of these Cyperaceae and *Ananas* paralogs is poorly supported. While the precise timing remains uncertain, the duplication that created the *NTT* and *NTT-like* clades predates the origin of the grass family and may correspond to the sigma duplication event early in the Poales (*60*). Prior to the diversification of the grass family, there was a second duplication event creating the *NTT-like1* and *NTT-like2* clades. The paralogous *NTT-like1* and *NTT-like2* genes are present in all sampled grasses and the timing of this duplication is consistent with the well-documented rho whole genome duplication event (*60*). While our phylogeny does not show support for *Ananas* or Cyperaceae paralogs in the *NTT* lineages, given the known history of whole genome duplications at the base of the grasses and Poales, it is more likely that a single duplication created paralogs of *NTT* and *NTT-like* in all Poales including *Ananas* and Cyperaceae rather than a more complicated series of duplication and loss consistent with the poorly supported topology we recovered.

While *tsh1* and orthologous *NTT* genes in the grass family have a conserved role in bract suppression, the duplication that created the *NTT* lineage clearly predates the origin of bracts, raising the question of what this clade of genes did before they were involved in bract suppression. In order to infer the likely ancestral expression of the *NTT* lineage we performed *in situ* hybridization using the likely *NTT* ortholog from *Cyperus*. The *Cyperus* inflorescence has prominent bracts and prophylls associated with all inflorescence branching events (Fig. 5B and C). *Cyperus NTT* RNA was present in a distinct boundary domain separating the spikelet meristem from lateral florets (Fig. 5D), but was not inbracts or other lateral organs. This boundary domain expression is similar to that reported for *HAN* in arabidopsis (*59*), and suggests that suppressed bract expression of *NTT* genes in the grass family is a neofunctionalization that arose after the duplication event that created the *NTT* lineage.

Neofunctionalization can involve changes to gene expression domains as well as to protein function. While grass *NTT* genes likely evolved a novel expression domain in the suppressed bract, it is not clear if the protein function also diverged. We reasoned that if both HAN and TSH1 proteins maintain an ancestral boundary domain function to suppress organ growth (*46, 47*), ectopic expression of either protein in young lateral primordia would suppress growth. Consequently, we ectopically expressed TSH1 and HAN in lateral organs of arabidopsis (Fig. 5E-5G). In order to avoid the deleterious effects of suppressing all leaf growth, we used the arabidopsis *AP3* promoter:LhG4 fusion (p*AP3*) to drive expression of 10-OP:*HAN* and 10-OP:*tsh1* cDNA fusions just in petal and stamen lateral primordia. Both p*AP3>>HAN* and p*AP3>>tsh1* flowers lacked petals and stamens, consistent with our hypothesis that HAN and TSH1 have a conserved boundary domain function of inhibiting organ growth. Intriguingly, the overexpression phenotype was not limited to petal and stamen suppression, but included growth of unorganized callus-like tissue in the same position of stamens and petals. This result was unexpected and suggests that *HAN* and *tsh1* are not only sufficient to inhibit growth, but also to promote de-differentiation and callus formation, which may be a result of the strong expression driven by the *AP3* promote combined with the apparent hormone modifying activity of *HAN (19–21)* and *tsh1* (this study).

The shift in *NTT* expression from boundary regions in a grass outgroup to the suppressed bract in the grasses likely results from grass *NTT* genes coming under the regulation of novel upstream factors. Since *tsh4* in maize and other grasses maintains an ancestral expression pattern in lateral organs (*5, 61, 62*), one possibility is that TSH4 binding to the *tsh1* promoter recruited this gene to a novel domain of inflorescence bracts. We examined 5 kb of 5’ promoter regions of *tsh1* and *NTT* genes in grasses and close outgroups *Joinvillea* and *Carex* to look for potential evidence of changes in TSH4 binding (Fig. 5H). We identified two syntenous conserved non-coding sequences (CNSs) in all promoters, with the exception of *Carex* which lacked one of the CNSs. We also mapped the distribution of the consensus SBP binding sites (GTAC; (*63*)), and found they were distributed randomly throughout all promoters. In addition, we observed a marked cluster of potential SBP binding sites in some promoters. In maize *tsh1*, this cluster overlapped with the known binding site of TSH4. A similar cluster was identified in all the core grass *NTT* promoters, but reduced (*Pharus* and *Joinvillea*) or lacking (*Carex*) outside the core grasses. While the relevance of the apparent gain in TSH4 binding site in grass *NTT* genes to *in vivo* binding dynamics will require further confirmation, the pattern we see is consistent with recruitment of *tsh1* by *tsh4* early in the evolution of the grass family.

Taken together, these results are consistent with a model in which a gene duplication event created the *NTT* lineage, which maintained the ancestral boundary domain function and expression. Later, the *NTT* lineage was recruited, possibly by *tsh4*, to the bract where it maintained its ancestral boundary domain protein function leading to inhibition of bract growth (Fig. 6).

**Fig. 6.**
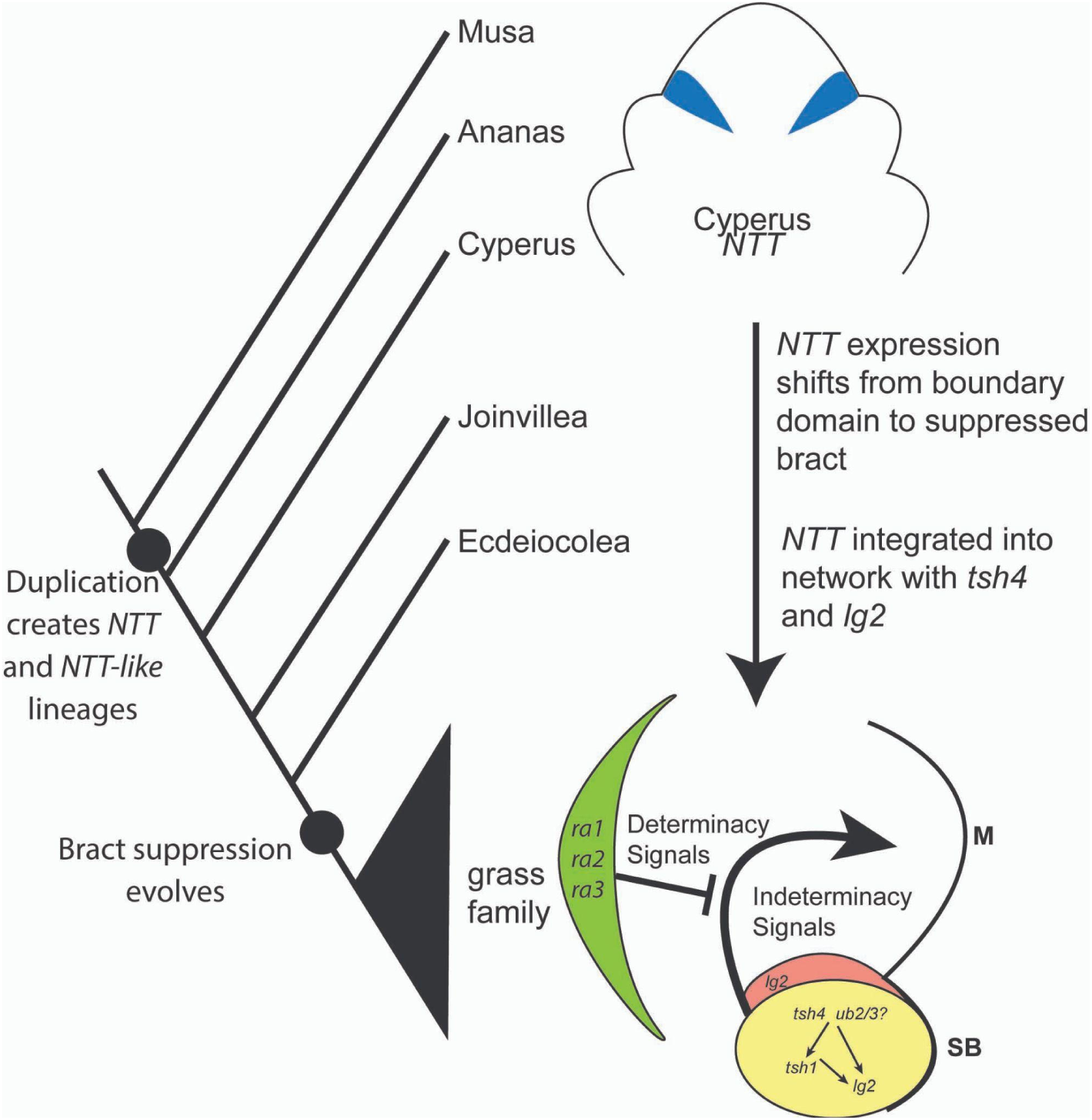
Proposed evolutionary origin and core network of bract suppression in the grass family. SB, suppressed bract. M, meristem.

## Discussion

We identified a core network composed of *tsh1, tsh4* and *lg2* that redundantly regulate bract suppression and branch meristem determinacy. This network is hierarchical with *tsh4* upstream of *tsh1*, and both *tsh4* and *tsh1* upstream of *lg2* (Fig. 6). *tsh1* and associated *NTT* genes in the grass family likely neofunctionalized, shifting an ancestral boundary domain gene to the bracts of the grass inflorescence, where they inhibit bract growth and promote branch indeterminacy (Fig. 6). These results provide novel insights into the regulatory structure of the bract suppression network, its evolutionary origin, and possible roles for bract suppression in grass inflorescence architecture.

### *tsh4* and *ub2*/*3* are master regulators of bract suppression associated with phase change

The transition from vegetative to reproductive development in grasses involves striking changes to plant development including a change in phyllotaxy, suppression of internode elongation, inhibition of leaf (bract) growth and associated promotion of branch meristem growth. The reproductive transition is the final in a series of phase changes that are regulated by a developmental mechanism that appears to be largely conserved across the angiosperms. This mechanism involves competing sets of transcription factors, each in turn regulated by microRNAs. The adult and floral phases are promoted by miR172 regulation of its AP2 family targets, while the juvenile phase is promoted by miR156 regulation of its SPL targets (*64*). Of the many morphological changes that occur during the transition to flowering, the miR156 targets *tsh4* and the paralogous *ub2* and *ub3* regulate bract suppression, tassel branching and meristem size (*7*). Thus, *SPL* genes are largely conserved in their phase transition roles, while their downstream targets can change in accordance with lineage specific developmental differences involved in floral transition such as bract suppression. Our data suggest that *tsh1* and *lg2* are downstream targets of *tsh4* in the maize, and possibly throughout the grass family. Given the residual expression of *tsh1* in *tsh4* mutants and the genetic redundancy of *ub2* and *ub3* with *tsh4* in bract suppression (*7*), *ub2* and *ub3* are strong candidates for promoting *tsh1* expression in parallel with *tsh4*. Our results are consistent with a model in which bract suppression in the grasses came under the control of *tsh4*/*ub2*/*ub3* as part of the reproductive transition placing *tsh4* at the top of a regulatory hierarchy of bract suppression and the associated promotion of inflorescence meristem indeterminacy (Fig. 6). This contrasts with the convergent suppression of bracts in arabidopsis, which is regulated by distinct genetic mechanisms with a large role played by the floral meristem identity gene *LFY*.

### *tsh1*/*NTT* genes in the grass family likely maintain boundary domain functions

Our data suggest that the novel function of *tsh1/NTT* genes in grass bract suppression evolved from an ancestral role in boundary domain promotion. Despite the evolution of a novel expression domain in the suppressed bract, TSH1 appears to have largely maintained the molecular activity of a boundary domain gene. While boundary domain genes include a diverse set of transcription factors, they share some molecular and morphogenetic properties including the ability to inhibit cell division and expansion in organ boundaries, direct growth of adjacent tissues, and regulate hormone homeostasis (*46, 65, 66*). Thus it appears that boundary domains simultaneously function as growth repressors and non-autonomous signaling centers, roles that evolved convergently for boundary domains in animals (*67*). Transcript profiling of *tsh1* in maize and ectopic expression of TSH1 in arabidopsis are consistent with a model in which TSH1 maintains ancestral boundary domain activities, but transferred them to a lateral organ primordium (bract) suppressing its growth and simultaneously promoting growth in the adjacent branch meristem.

### Developmental regulation of ligules, bracts, and branch meristem determinacy is tightly correlated in the grasses

Previous work in maize has shown a recurring pleiotropy in which ligule mutants also have tassel branching defects (*48, 49*). Here we show that, at least for *lg2*, this pleiotropy extends to bract suppression as *lg2* interacts with both *tsh1* and *tsh4* to regulate both branch meristem determinacy and bract suppression. Given the *tsh1 lg2* interaction in inflorescence development, it is interesting to note that *tsh1* expression is localized to the ligule in both maize and rice (*50, 51*), although no obvious functional role for *tsh1* in ligule development has yet emerged. It is not immediately clear why bract suppression and inflorescence branching would be under the control of the same genes that are necessary for ligule establishment. Insofar as the ligule can be understood as a boundary that separates the sheath and blade in grass leaves, the correlation of ligule development with bract growth and inflorescence branching further underscores the important role of boundary domain genes in these distinct developmental contexts. While the suppressed bract is a novel trait in the grass family, the origin of ligules is less clear (*68*). Future work exploring the intersection of ligule development and bract suppression in an expanded phylogenetic context may shed light on the molecular and evolutionary mechanisms by which these traits arose and became integrated during grass inflorescence branching.

### Bract suppression may be an indirect effect of creating a branch meristem indeterminacy promoting region acting in opposition *ramosa* genes

Inflorescence architecture is highly diverse across the angiosperms, and particularly in the grass family. Establishing this architecture requires regulation of meristem determinacy during ontogeny of the inflorescence. After initiation, lateral meristems can either continue growing and branching (indeterminacy) or form a limited number of floral primordia before the meristem is consumed (determinacy)(*26*).

The size and activity of the meristem is coordinated by complex interactions of non-autonomous factors that signal from multiple domains within and around the meristem. Meristem growth vs. non-growth is not a simple switch, but a complex read-out of competing signals that either promote or inhibit meristem size (*69*), including signals originating outside the meristem proper from adjacent lateral organs (*70, 71*). While many aspects of the meristem growth network are likely common to all meristems in the plant, how these meristem dynamics are regulated in different developmental contexts to alter plant architecture is less clear. Maize inflorescence architecture mutants suggest that determinacy of branch meristems in the inflorescence involve additional regions of non-autonomous signaling specific to the grass family.

The *ramosa* (*ra*) genes, *ra1, ra2*, and *ra3*, are core regulators of branch meristem determinacy in maize. However, *ra* genes are not expressed in the meristem itself, but rather in a boundary domain adaxial to the meristem (*72*–*74*) that *ra* genes establish a determinacy signaling center adjacent to the meristem. Our work demonstrates that *tsh1, tsh4* and *lg2* interact in a network necessary not only for bract suppression, but also for meristem indeterminacy, thus functioning in opposition to *ramosa* genes. Supporting this, *tsh4* is epistatic to *ra* mutants with respect to inflorescence branch determinacy (*5*). Thus genetic evidence suggests that the indeterminacy promoting effects of *tsh4* are negatively regulated *ra* signaling. Furthermore, *tsh1, tsh4* and *lg2* are not expressed directly in the meristem, but act from an adjacent domain, analogous to *ra* genes but in the meristem subtending bract rather than adaxial to the meristem. Inflorescence meristem branching in maize thus appears to be under the control of antagonistic signaling centers loosely analogous to the interaction of signaling domains that internally regulate meristem size (*26, 75*).

While the evidence presented here supports a signaling role for *tsh* genes in the promotion of branch determinacy, the nature of the non-autonomous signal originating in the bract is still uncertain. Mobile biomolecules including proteins, small peptides, RNA, hormones or other small molecules are all possible. An intriguing possibility is that the extensive diversification of branching architecture in the grasses was facilitated by the integration of indeterminacy and determinacy signals emanating from the suppressed bract and *ramosa* genes respectively to create a grass-specific mechanism to regulate inflorescence branching. Future work to elucidate the maize bract indeterminacy signal and its interaction with antagonistic determinacy signals will provide a framework for understanding the developmental constraints regulating inflorescence architecture in this agronomically important species.

## Materials and Methods

### Plant materials

Two new alleles of *tsh4* were isolated from distinct sources. *tsh4-rm* (originally *tsh2*) was identified in a *Mutator*-transposon active population, and was introgressed over 5 times to the reference B73 background prior to all experiments described here. In a screen for genetic modifiers of *tsh1-2* (A619 background), a phenotypically weak allele, we identified several *enhancers of tasselsheath1* (*ent*) mutants including one we designated *ent*-355*, that was subsequently re-named *tsh4-ent*-355* based on mapping and allelism with *tsh4-ref. lg2-R* was introgressed 4 times into B73 before generating *lg2; tsh1*-*ref* and *lg2; tsh4-rm* double mutant populations. B73, *tsh1-ref, tsh4-rm*, and *tsh1-ref tsh4-rm* mutants used for LM-RNA-seq assays were grown in 5-gallon pots in a greenhouse at 24 °C with supplemental lights for 16-h light/ 8-h dark period.

### Generating double mutants, genotyping and phenotypic analysis

*tsh1*-*ref* (B73) was crossed as a female to *tsh4*-*rm* (B73) to generate a *tsh1*/*+, tsh4*/*+* segregating population. *lg2*-*R* (B73) was crossed as female to *tsh1*-*ref* and *tsh4*-*rm* to generate *tsh1*/*+, lg2*/*+* and *tsh4*/*+, lg2*/*+* segregating populations, respectively. Each segregating population was grown at an irrigated field in Spanish Fork, Utah and genotyped by PCR using NEB OneTaq DNA polymerase and (see table S13 for primer sequence and genotyping instructions). Mature tassels from individual plants were collected and tassel-related phenotypes were inspected manually.

### Scanning electron microscopy

Dissected ear and tassel primordia were fixed overnight in FAA (4% formalin, 50% ethanol, 5% acetic acid, 0.01% Triton X-100), dehydrated through an ethanol series, and transitioned to 100% acetone before drying using a 931.GL Supercritical Autosamdri critical point dryer (CPD) (Tousimis, Maryland) according to the manufacturer’s protocol. Dried samples were mounted on stubs and sputter coated with gold:palladium (80:20) with a Quorum Q150TES (Quorum Technologies, East Sussex, England), before imaging on a XL30 FEI Scanning Electyron Microscope (TSS Microscopy, Hillsboro, OR, USA) with an acceleration voltage of 5-30 kV under high vacuum mode (<20 mBar) with a working distance of 10 to 20 mm.

### Laser microdissection and RNA-seq library preparation

LM-seq was performed largely as described by (*76*). Briefly, ear primordia were fixed overnight in 3.5% paraformaldehyde, dehydrated through an ethanol series, transferred to Histoclear (National Diagnostics) before embedding in paraffin. Embedded samples were sectioned at 5 µm and mounted onto charged HistoBond slides (VWR International). The slides were then subjected to laser microdissection by using a PALM microbeam system (Zeiss). Three biological replicates were prepared per genotype (B73, *tsh1, tsh4*, and *tsh1 tsh4*). Approximately 750 microns of cells were harvested for each biological replicate. RNA was extracted from microdissected tissues with the PicoPure RNA Isolation Kit (Life Technologies, Carlsbad, CA) and in vitro amplified using a TargetAmp 2-round aRNA Amplification Kit 2.0 (Epicentre, Madison, WI). RNA-seq libraries were constructed using a TruSeq Stranded mRNA Library Prep Kit and quantified on an Agilent bioanalyzer (Agilent) and single-end 125bp sequences generated on an Illumina Hi-seq 2500.

### Differential expression analysis of RNA-seq data

Differential expression analysis was performed as previously described (*77*) with minor modification. Twelve RNA-seq libraries were sequenced and a total of ∼958 million single-end (SE) raw reads were obtained with an average of 79.9 million reads per library. The overall quality of our sequencing data was assessed using FastQC and the raw reads were filtered using Trimmomatic v.0.36 to trim and remove low-quality reads and adapter sequences. The filtered reads were mapped to the maize B73 reference genome version 3, release 31 (AGPv3.31) using STAR aligner v.2.6.0a with default parameter settings. Total mapped and uniquely mapped reads are summarized in table S14. A read count matrix including all samples was generated by aggregating the raw counts of mapped reads for a given gene in each sample using featureCounts with reference to 39479 maize gene models in AGPv3.31. The read count matrix was subjected to differential gene expression analysis using a Bioconductor R package edgeR v.3.22.5. Briefly, genes with ubiquitously low expression were filtered out from the read count matrix in order to improve differential expressed gene detection sensitivity and only the genes that had count-per-million (CPM) value >0.25 in at least three libraries were retained. This resulted in a filtered read count matrix containing 20168 expressed genes in the samples (table S1). The filtered read count matrix was normalized for compositional bias between libraries using a trimmed means of M values (TMM) method and then used to detect genes with differential expression between pairwise samples. Genes with an adjusted p-value (q-value) <= 0.05 and an absolute value of log2 fold changes (FC) >=1 were considered as differentially expressed.

### Gene co-expression cluster analysis

Co-expression analysis was performed as previously described (*77*) with minor modification. The 1389 genes that were differentially expressed between the wildtype (B73) and the three *tsh* mutants (*tsh1, tsh4* and *tsh1*;*tsh4*) were subjected to co-expression cluster analysis across all samples using a Bioconductor R package coseq v1.5.2. The raw read count matrix of the 1389 genes in the 12 RNA-seq libraries were converted into a RPKM (Reads Per Kilobase of transcript, per Million mapped reads) matrix (table S15) which was then used as an input in coseq for co-expression analysis. Briefly, log CLR-transformation and TMM normalization were applied to the gene expression matrix to normalize the expression of genes and the K-means algorithm was used to identify the co-expressed clusters across all samples. A range of clusters from 2 to 20 was tested to identify the optimal number of clusters. The K-means algorithm embedded in the coseq() function was repeated for 40 iterations (counts, K=2:20, transformation=“logclr”,norm=“TMM”, model=“kmeans”) and the resulting number of clusters in each run was recorded. The most frequently occurring number of clusters was selected as the optimal number of clusters, and genes that were assigned to these clusters were retained for cluster visualization and gene ontology enrichment analysis.

### Gene ontology enrichment analysis

Statistically enriched (overrepresented with an adjusted p-value <= 0.05) gene ontology (GO) terms for genes differentially expressed between pairwise samples or for genes assigned to certain co-expression clusters were identified using Singular Enrichment Analysis (SEA) in AgriGO v2.0 at http://systemsbiology.cau.edu.cn/agriGOv2/ with default parameter settings. After collapsing and removing redundant or very high-level terms, the most statistically enriched GO terms were plotted in ggplot2 for visualization.

### TSH4 ChIP-PCR

Maize B73 plants were grown in the experimental field of the Plant Gene Expression Center, UC Berkeley. Young ear primordia smaller than 5 mm were carefully dissected. About 1 g of tissue per biological replicate was fixed in 1% formaldehyde solution for 10 min under vacuum, and quenched by adding glycine to a final concentration of 0.1 M. Nuclei extraction and chromatin immunoprecipitation using TSH4 antibody were performed as described previously (*77*). Normal goat anti-rabbit IgG was used as a negative control. To validate the putative TSH4-binding targets, three biological replicates of immunoprecipitated DNA in ChIP were applied for each qPCR using respective primer pairs listed in table S13 with Fast Evagreen qPCR mix. Relative enrichment was calculated using the ΔCt (threshold cycle) method, and significant difference was evaluated through t-test between anti-TSH4 ChIPed samples and IgG control.

### LG2 Antibody Generation and Immunolocalization

Full length LG2 coding sequence was cloned into gateway vector pDEST17. N terminal HIS tagged full length LG2 was expressed in rosetta cells and purified in 8M Urea. The antibodies were produced in guinea pigs (Cocalico Biologicals). Whole serum was tested for reactivity via dot blot and then purified first against HIS protein, then against the N terminus of LG2 (residues: 1-200, cloned into pDEST15, produced in rosetta cells) fused to GST as described by (*5*). Specificity was tested using immunolocalization and western blot in wild-type and *lg2* tissue. Immunolocalization used a protocol based on (*78*), and was as follows. Slides were deparaffinized using histoclear, then rehydrated through an ethanol series to water. Samples were then boiled in 10mM sodium citrate, pH6, for 10 minutes to retrieve the antigens. Blocking was carried out in 1% BSA/PBS/0.3% Triton-X100 for 3 hours. Slides were incubated overnight in the primary antibody, before washing in PBS/0.3% Triton-X100. They were then incubated with secondary antibody (goat anti-guinea pig alkaline phosphatase conjugate (Bethyl, #A60-110AP)) at room-temperature for 2 hours. Slides were then were incubated in a 1:50 dilution of NBT/BCIP (Roche, #11681451001) in 0.05M MgCl_2_/TBS, pH9.5 until a dark precipitate was observed. These slides were then imaged on a Leica MZ16-F dissecting microscope with an attached canon EOS 250D camera, in water. LG2 antibody was used at a 1:300 dilution, and goat anti-guinea pig alkaline phosphatase was used at a 1:400 dilution in 1% bovine serum albumin (BSA) in PBS.

### Isolation of *NTT* orthologs from the Poales and Phylogenetic Analysis

Genomic DNA and/or total RNA were isolated from young inflorescences of *Hyparrhenia hirta, Streptochaeta angustifolia, Pharus latifolia, Joinvillea ascendens, Elegia tectorum, Georgeantha hexandra*, and *Cyperus papyrus*. cDNA was generated using the Superscript III First-Strand kit (Thermo Fisher) according to the manufacturer’s protocol with a modified polyT primer (table S13). A series of degenerate primers (table S13) designed for the HAN domain or the GATA domain were used in combination with a polyT primer to isolate the 3’ sequence using 3’ RACE, while primers designed to conserved noncoding sequences in the 5’ promoter with gene specific reverse primers were used to isolate the 5’ end of genes where possible. These amplified sequences were aligned using CLUSTALX, and the resulting alignment adjusted by hand using MacClade to create a preliminary alignment. This preliminary alignment was used as the prior to search the Gramene and Phytozome coding sequence databases using a hidden Markov model (HMM) in HMMER v.3.1b2 to isolate *NTT/HAN* orthologs from *Arabidopsis thaliana, Populus trichocarpa, Selaginella moellendorffii, Physcomitrella patens, Marchantia polymorpha, Amborella trichopoda, Zea mays, Miscanthus sinensis, Sorghum bicolor, Setaria italica, Panicum virgatum, Brachypodium distachyon, Hordeum vulgare, Triticum aestivum, Oryza sativa, Musa acuminata, Dioscorea rotunda, Ananas comosus*.

All identified *NTT/HAN* orthologs were aligned using MAFFT v.7.313, which was then filtered for homoplastic positions by Noisy v.1.5.12. Finally, the alignment was tested for the best substitution model and used to infer a maximum-likelihood gene tree and 1000 bootstrap replicates using IQTree v.1.6.3. The best-fit model selected by IQTree was TVM+F+R3. The tree was visualized using R v.4.0.2., and a subclade containing the gene of interest (maize TSH1) out to the nearest outgroup clade (containing genes from *S. moellendorffii, P. patens*, and *M. polymorpha*) was selected for further refinement. The process described above was repeated, but using only the genes included in the subclade as an input to MAFFT. Finally, *tsh1* orthologous sequences identified from the recently published *Cyperus littledalei* genome (*79*) were manually aligned to the final alignment using Mesquite v.3.61, and this alignment was used to create the tree using IQTree as described above.

### In situ hybridization

An anti-sense T7 probe, labelled with dig-UTP (Roche) was synthesized for the full-length cDNA for *CyperusNTT* using the Invitrogen SuperscriptIII kit according to the manufacturer’s protocol. Tissue preparation and *in situ* hybridization was performed as previously described (*3*).

### Transgenic Arabidopsis

Coding sequence for *tsh1* and *HAN* were amplified from cDNA using primers (table S13) with 5’ *XhoI* (5’) and 3’ *BamHI* (3’) sites and cloned into the pBJ36 vector downstream of the 10-OP promoter. *NotI* fragments containing the 10-OP:*tsh1* and 10-OP:*HAN* gene promoter fusions were then subcloned from pBJ36 into the pMLBART27 binary vector and subsequently transformed into *Agrobacterium* and used to transform wild type *Arabidopsis thaliana* Col.. Transgenic lines containing 10-OP:*tsh1* and 10-OP:*HAN* were crossed to the pAP3:Lhg4 driver line (*70*).

### Promoter Analysis

Genomic sequence containing 5kb of 5’ promoter region upstream of the start codon for *tsh1* (*Zea mays*), sorghum *NTT* (*Sorghum bicolor*), setaria *NTT* (*Setaria italica*), *NL1* (*Oryza sativa*), brachypodium *NTT* (*Brachypodium distachyon*), *Pharus latifolia NTT, Joinvillea ascendens NTT*, and *Carex littledalei NTT* was isolated the respective published genomes of each species. Pairwise alignments of all promoters were performed with blastn to identify significant stretches of nucleotide identity spanning at least 15 bp. After aligning all pairs of sequences, two regions of similarity from blastn alignments were identified with only *Carex* lacking one of these. Since they were syntenously arranged in all promoters we designated these regions conserved non-coding sequences.

## Acknowledgments

We thank Michael Lewis for providing images of *lg2* inflorescences and preliminary analysis of LG2 localization, as well as Josh Strable for his helpful feedback on the manuscript. We sincerely apologize to colleagues whose work could not be cited here due to the space limitation.

## Funding

This work was supported by a NSF CAREER award to CJW (IOS-1253421).

## Author contributions

C.W. conceived and supervised the project. Y.G.X., J.Y.G., Z.B.D., A.R., S.M., S.B., E.P., E.B., M.B., G.C., A.L.E. and C.W. performed experiments. M.J.C. contributed materials and reagents for LM RNA-seq sample preparation. Y.G.X. and C.W. analyzed the data and wrote the original manuscript with the input from all authors. All authors have read, edited and approved the paper.

## Competing interests

The authors declare that they have no competing interests.

## Data and materials availability

All raw LM-RNA-seq data generated in this study have been deposited in the National Center for Biotechnology Information Gene Expression Omnibus, www.ncbi.nlm.nih.gov/ (accession number GSE179997). All data needed to evaluate the conclusions in the paper are present in the paper and/or the Supplementary Materials.

## Notes

### Competing Interest Statement

The authors have declared no competing interest.

## References

1. F. Weberling, Morphology of Flowers and Inflorescences (Cambridge University Press, Cambridge, 1989).

2. M. K. Ritter, C. M. Padilla, R. J. Schmidt, The maize mutant barren stalk1 is defective in axillary meristem development. Am. J. Bot. 89, 203–210 (2002).

3. C. J. Whipple, D. H. Hall, S. DeBlasio, F. Taguchi-Shiobara, R. J. Schmidt, D. P. Jackson, A conserved mechanism of bract suppression in the grass family. Plant Cell. 22, 565–578 (2010).

4. I. A. Al-Shehbaz, M. A. Beilstein, E. A. Kellogg, Systematics and phylogeny of the Brassicaceae (Cruciferae): an overview. Osterr. bot. Z. 259, 89–120 (2006).

5. G. Chuck, C. Whipple, D. Jackson, S. Hake, The maize SBP-box transcription factor encoded by tasselsheath4 regulates bract development and the establishment of meristem boundaries. Development. 137, 1243–1250 (2010).

6. G. Chuck, A. M. Cigan, K. Saeteurn, S. Hake, The heterochronic maize mutant Corngrass1 results from overexpression of a tandem microRNA. Nat. Genet. 39, 544–549 (2007).

7. G. S. Chuck, P. J. Brown, R. Meeley, S. Hake, Maize SBP-box transcription factors unbranched2 and unbranched3 affect yield traits by regulating the rate of lateral primordia initiation. Proc. Natl. Acad. Sci. U. S. A. 111, 18775–18780 (2014).

8. D. Weigel, J. Alvarez, D. R. Smyth, M. F. Yanofsky, E. M. Meyerowitz, LEAFY controls floral meristem identity in Arabidopsis. Cell. 69, 843–859 (1992).

9. S. R. Hepworth, Y. Zhang, S. McKim, X. Li, G. W. Haughn, BLADE-ON-PETIOLE–Dependent Signaling Controls Leaf and Floral Patterning in Arabidopsis. Plant Cell. 17, 1434–1448 (2005).

10. M. Norberg, M. Holmlund, O. Nilsson, The BLADE ON PETIOLE genes act redundantly to control the growth and development of lateral organs. Development. 132, 2203–2213 (2005).

11. K. D. Allen, I. M. Sussex, Falsiflora and anantha control early stages of floral meristem development in tomato (Lycopersicon esculentum Mill.). Planta. 200, 254–264 (1996).

12. N. Molinero-Rosales, M. Jamilena, S. Zurita, P. Gómez, J. Capel, R. Lozano, FALSIFLORA, the tomato orthologue of FLORICAULA and LEAFY, controls flowering time and floral meristem identity. Plant J. 20, 685–693 (1999).

13. J. Hofer, L. Turner, R. Hellens, M. Ambrose, P. Matthews, A. Michael, N. Ellis, UNIFOLIATA regulates leaf and flower morphogenesis in pea. Curr. Biol. 7, 581–587 (1997).

14. K. Ikeda-Kawakatsu, M. Maekawa, T. Izawa, J.-I. Itoh, Y. Nagato, ABERRANT PANICLE ORGANIZATION 2/RFL, the rice ortholog of Arabidopsis LEAFY, suppresses the transition from inflorescence meristem to floral meristem through interaction with APO1. Plant J. 69, 168–180 (2012).

15. K. Bomblies, R.-L. Wang, B. A. Ambrose, R. J. Schmidt, R. B. Meeley, J. Doebley, Duplicate FLORICAULA/LEAFY homologs zfl1 and zfl2 control inflorescence architecture and flower patterning in maize. Development. 130, 2385–2395 (2003).

16. Z. Dong, W. Li, E. Unger-Wallace, J. Yang, E. Vollbrecht, G. Chuck, Ideal crop plant architecture is mediated by tassels replace upper ears1, a BTB/POZ ankyrin repeat gene directly targeted by TEOSINTE BRANCHED1. Proc. Natl. Acad. Sci. U. S. A. 114, E8656–E8664 (2017).

17. M. Xu, T. Hu, J. Zhao, M.-Y. Park, K. W. Earley, G. Wu, L. Yang, R. S. Poethig, Developmental Functions of miR156-Regulated SQUAMOSA PROMOTER BINDING PROTEIN-LIKE (SPL) Genes in Arabidopsis thaliana. PLoS Genet. 12, e1006263 (2016).

18. S. Schwarz, A. V. Grande, N. Bujdoso, H. Saedler, P. Huijser, The microRNA regulated SBP-box genes SPL9 and SPL15 control shoot maturation in Arabidopsis. Plant Mol. Biol. 67, 183–195 (2008).

19. T. Nawy, M. Bayer, J. Mravec, J. Friml, K. D. Birnbaum, W. Lukowitz, The GATA factor HANABA TARANU is required to position the proembryo boundary in the early Arabidopsis embryo. Dev. Cell. 19, 103–113 (2010).

20. L. Ding, S. Yan, L. Jiang, W. Zhao, K. Ning, J. Zhao, X. Liu, J. Zhang, Q. Wang, X. Zhang, HANABA TARANU (HAN) bridges meristem and organ primordia boundaries through PINHEAD, JAGGED, BLADE-ON-PETIOLE2 and CYTOKININ OXIDASE 3 during flower development in Arabidopsis. PLoS Genet. 11, e1005479 (2015).

21. X. Zhang, Y. Zhou, L. Ding, Z. Wu, R. Liu, Transcription repressor HANABA TARANU controls flower development by integrating the actions of multiple hormones, floral organ specification genes, and GATA3 …. The Plant (2013) (available at http://www.plantcell.org/content/25/1/83.short).

22. E. S. Coen, J. M. Nugent, Evolution of flowers and inflorescences. Development. 1994, 107–116 (1994).

23. G. Chuck, E. Bortiri, The unique relationship between tsh4 and ra2 in patterning floral phytomers. Plant Signal. Behav. 5, 979–981 (2010).

24. J. W. Chandler, Floral meristem initiation and emergence in plants. Cell. Mol. Life Sci. 69, 3807– 3818 (2012).

25. D. A. Baum, C. D. Day, Cryptic bracts exposed: insights into the regulation of leaf expansion. Dev. Cell. 6, 318–319 (2004).

26. C. J. Whipple, Grass inflorescence architecture and evolution: the origin of novel signaling centers. New Phytol. 216, 367–372 (2017).

27. D. Jackson, B. Veit, S. Hake, Expression of maize KNOTTED1 related homeobox genes in the shoot apical meristem predicts patterns of morphogenesis in the vegetative shoot. Development. 120, 405– 413 (1994).

28. J. Strable, J. G. Wallace, E. Unger-Wallace, S. Briggs, P. J. Bradbury, E. S. Buckler, E. Vollbrecht, Maize YABBY Genes drooping leaf1 and drooping leaf2 Regulate Plant Architecture. The Plant Cell. 29 (2017), pp. 1622–1641.

29. R. Sarojam, P. G. Sappl, A. Goldshmidt, I. Efroni, S. K. Floyd, Y. Eshed, J. L. Bowman, Differentiating Arabidopsis Shoots from Leaves by Combined YABBY Activities. The Plant Cell. 22 (2010), pp. 2113–2130.

30. J.A. Aguilar-Martínez, N. Sinha, Analysis of the role of Arabidopsis class I TCP genes AtTCP7, AtTCP8, AtTCP22, and AtTCP23 in leaf development. Front. Plant Sci. 4, 406 (2013).

31. J. F. Palatnik, E. Allen, X. Wu, C. Schommer, R. Schwab, J. C. Carrington, D. Weigel, Control of leaf morphogenesis by microRNAs. Nature. 425, 257–263 (2003).

32. P. Ballester, M. Navarrete-Gómez, P. Carbonero, L. Oñate-Sánchez, C. Ferrándiz, Leaf expansion in Arabidopsis is controlled by a TCP-NGA regulatory module likely conserved in distantly related species. Physiol. Plant. 155, 21–32 (2015).

33. J. H. Kim, D. Choi, H. Kende, The AtGRF family of putative transcription factors is involved in leaf and cotyledon growth in Arabidopsis. Plant J. 36, 94–104 (2003).

34. H. Nelissen, D. Eeckhout, K. Demuynck, G. Persiau, A. Walton, M. van Bel, M. Vervoort, J. Candaele, J. De Block, S. Aesaert, M. Van Lijsebettens, S. Goormachtig, K. Vandepoele, J. Van Leene, M. Muszynski, K. Gevaert, D. Inzé, G. De Jaeger, Dynamic Changes in ANGUSTIFOLIA3 Complex Composition Reveal a Growth Regulatory Mechanism in the Maize Leaf. The Plant Cell. 27 (2015), pp. 1605–1619.

35. A. Gallavotti, Q. Zhao, J. Kyozuka, R. B. Meeley, M. K. Ritter, J. F. Doebley, M.E. Pè, R. J. Schmidt, The role of barren stalk1 in the architecture of maize. Nature. 432, 630–635 (2004).

36. A. Gallavotti, S. Malcomber, C. Gaines, S. Stanfield, C. Whipple, E. Kellogg, R. J. Schmidt, BARREN STALK FASTIGIATE1 is an AT-hook protein required for the formation of maize ears. Plant Cell. 23, 1756–1771 (2011).

37. M. P. Joseph, C. Papdi, L. Kozma-Bognár, I. Nagy, M. López-Carbonell, G. Rigó, C. Koncz, L. Szabados, The Arabidopsis ZINC FINGER PROTEIN3 Interferes with Abscisic Acid and Light Signaling in Seed Germination and Plant Development. Plant Physiol. 165, 1203–1220 (2014).

38. Y. Hu, X. Han, M. Yang, M. Zhang, J. Pan, D. Yu, The Transcription Factor INDUCER OF CBF EXPRESSION1 Interacts with ABSCISIC ACID INSENSITIVE5 and DELLA Proteins to Fine-Tune Abscisic Acid Signaling during Seed Germination in Arabidopsis. Plant Cell. 31, 1520–1538 (2019).

39. K. Chen, G.-J. Li, R. A. Bressan, C.-P. Song, J.-K. Zhu, Y. Zhao, Abscisic acid dynamics, signaling, and functions in plants. J. Integr. Plant Biol. 62, 25–54 (2020).

40. M. G. Heisler, M. E. Byrne, Progress in understanding the role of auxin in lateral organ development in plants. Curr. Opin. Plant Biol. 53, 73–79 (2020).

41. Y. Hu, Q. Xie, N.-H. Chua, The Arabidopsis auxin-inducible gene ARGOS controls lateral organ size. Plant Cell. 15, 1951–1961 (2003).

42. P. Hedden, V. Sponsel, A Century of Gibberellin Research. J. Plant Growth Regul. 34, 740–760 (2015).

43. J. Park, K. T. Nguyen, E. Park, J.-S. Jeon, G. Choi, DELLA Proteins and Their Interacting RING Finger Proteins Repress Gibberellin Responses by Binding to the Promoters of a Subset of Gibberellin-Responsive Genes in Arabidopsis. The Plant Cell. 25 (2013), pp. 927–943.

44. S. Gazzarrini, Y. Tsuchiya, S. Lumba, M. Okamoto, P. McCourt, The transcription factor FUSCA3 controls developmental timing in Arabidopsis through the hormones gibberellin and abscisic acid. Dev. Cell. 7, 373–385 (2004).

45. Z.-L. Zhang, M. Ogawa, C. M. Fleet, R. Zentella, J. Hu, J.-O. Heo, J. Lim, Y. Kamiya, S. Yamaguchi, T.-P. Sun, Scarecrow-like 3 promotes gibberellin signaling by antagonizing master growth repressor DELLA in Arabidopsis. Proc. Natl. Acad. Sci. U. S. A. 108, 2160–2165 (2011).

46. S. R. Hepworth, V. A. Pautot, Beyond the Divide: Boundaries for Patterning and Stem Cell Regulation in Plants. Front. Plant Sci. 6, 1052 (2015).

47. A. E. Richardson, S. Hake, Drawing a Line: Grasses and Boundaries. Plants. 8 (2018), doi:10.3390/plants8010004.

48. J. Walsh, M. Freeling, The liguleless2 gene of maize functions during the transition from the vegetative to the reproductive shoot apex. Plant J. 19, 489–495 (1999).

49. J. Moon, H. Candela, S. Hake, The Liguleless narrow mutation affects proximal-distal signaling and leaf growth. Development. 140, 405–412 (2013).

50. L. Wang, H. Yin, Q. Qian, J. Yang, C. Huang, X. Hu, D. Luo, NECK LEAF 1, a GATA type transcription factor, modulates organogenesis by regulating the expression of multiple regulatory genes during reproductive development in rice. Cell Res. 19, 598–611 (2009).

51. R. Johnston, M. Wang, Q. Sun, A. W. Sylvester, S. Hake, M. J. Scanlon, Transcriptomic analyses indicate that maize ligule development recapitulates gene expression patterns that occur during lateral organ initiation. Plant Cell. 26, 4718–4732 (2014).

52. K.-I. Hibara, M. R. Karim, S. Takada, K.-I. Taoka, M. Furutani, M. Aida, M. Tasaka, Arabidopsis CUP-SHAPED COTYLEDON3 regulates postembryonic shoot meristem and organ boundary formation. Plant Cell. 18, 2946–2957 (2006).

53. D.-K. Lee, M. Geisler, P. S. Springer, LATERAL ORGAN FUSION1 and LATERAL ORGAN FUSION2 function in lateral organ separation and axillary meristem formation in Arabidopsis. Development. 136, 2423–2432 (2009).

54. C. Gómez-Mena, R. Sablowski, ARABIDOPSIS THALIANA HOMEOBOX GENE1 establishes the basal boundaries of shoot organs and controls stem growth. Plant Cell. 20, 2059–2072 (2008).

55. M. A. Moreno, L. C. Harper, R. W. Krueger, S. L. Dellaporta, M. Freeling, liguleless1 encodes a nuclear-localized protein required for induction of ligules and auricles during maize leaf organogenesis. Genes Dev. 11, 616–628 (1997).

56. W. A. Ricci, Z. Lu, L. Ji, A. P. Marand, C. L. Ethridge, N. G. Murphy, J. M. Noshay, M. Galli, M.K. Mejía-Guerra, M. Colomé-Tatché, F. Johannes, M. J. Rowley, V. G. Corces, J. Zhai, M. J. Scanlon, E. S. Buckler, A. Gallavotti, N. M. Springer, R. J. Schmitz, X. Zhang, Widespread long-range cis-regulatory elements in the maize genome. Nat Plants. 5, 1237–1249 (2019).

57. J. Walsh, C. A. Waters, M. Freeling, The maize gene liguleless2 encodes a basic leucine zipper protein involved in the establishment of the leaf blade-sheath boundary. Genes & Development. 12 (1998), pp. 208–218.

58. K. Houston, A. Druka, N. Bonar, M. Macaulay, U. Lundqvist, J. Franckowiak, M. Morgante, N. Stein, R. Waugh, Analysis of the barley bract suppression gene Trd1. Theor. Appl. Genet. 125, 33–45 (2012).

59. Y. Zhao, L. Medrano, K. Ohashi, J. C. Fletcher, H. Yu, H. Sakai, E. M. Meyerowitz, HANABA TARANU is a GATA transcription factor that regulates shoot apical meristem and flower development in Arabidopsis. Plant Cell. 16, 2586–2600 (2004).

60. M. R. McKain, H. Tang, J. R. McNeal, S. Ayyampalayam, J. I. Davis, C. W. dePamphilis, T. J. Givnish, J. C. Pires, D. W. Stevenson, J. H. Leebens-Mack, A Phylogenomic Assessment of Ancient Polyploidy and Genome Evolution across the Poales. Genome Biol. Evol. 8, 1150–1164 (2016).

61. J.-W. Wang, R. Schwab, B. Czech, E. Mica, D. Weigel, Dual effects of miR156-targeted SPL genes and CYP78A5/KLUH on plastochron length and organ size in Arabidopsis thaliana. Plant Cell. 20, 1231–1243 (2008).

62. K. Miura, M. Ikeda, A. Matsubara, X.-J. Song, M. Ito, K. Asano, M. Matsuoka, H. Kitano, M. Ashikari, OsSPL14 promotes panicle branching and higher grain productivity in rice. Nat. Genet. 42, 545–549 (2010).

63. X. Liang, T. J. Nazarenus, J. M. Stone, Identification of a consensus DNA-binding site for the Arabidopsis thaliana SBP domain transcription factor, AtSPL14, and binding kinetics by surface plasmon resonance. Biochemistry. 47, 3645–3653 (2008).

64. C. Li, B. Zhang, MicroRNAs in Control of Plant Development. J. Cell. Physiol. 231, 303–313 (2016).

65. Q. Wang, A. Hasson, S. Rossmann, K. Theres, Divide et impera: boundaries shape the plant body and initiate new meristems. New Phytol. 209, 485–498 (2016).

66. A. Maugarny-Calès, M. Cortizo, B. Adroher, N. Borrega, B. Gonçalves, G. Brunoud, T. Vernoux, N. Arnaud, P. Laufs, Dissecting the pathways coordinating patterning and growth by plant boundary domains. PLoS Genet. 15, e1007913 (2019).

67. C. Kiecker, A. Lumsden, Compartments and their boundaries in vertebrate brain development. Nat. Rev. Neurosci. 6, 553–564 (2005).

68. P. A. Conklin, J. Strable, S. Li, M. J. Scanlon, On the mechanisms of development in monocot and eudicot leaves. New Phytol. 221, 706–724 (2019).

69. H. Han, X. Liu, Y. Zhou, Transcriptional circuits in control of shoot stem cell homeostasis. Curr. Opin. Plant Biol. 53, 50–56 (2020).

70. A. Goldshmidt, J. P. Alvarez, J. L. Bowman, Y. Eshed, Signals derived from YABBY gene activities in organ primordia regulate growth and partitioning of Arabidopsis shoot apical meristems. Plant Cell. 20, 1217–1230 (2008).

71. B. I. Je, J. Gruel, Y. K. Lee, P. Bommert, E. D. Arevalo, A. L. Eveland, Q. Wu, A. Goldshmidt, R. Meeley, M. Bartlett, M. Komatsu, H. Sakai, H. Jönsson, D. Jackson, Signaling from maize organ primordia via FASCIATED EAR3 regulates stem cell proliferation and yield traits. Nat. Genet. 48, 785–791 (2016).

72. E. Vollbrecht, P. S. Springer, L. Goh, E. S. Buckler 4th, R. Martienssen, Architecture of floral branch systems in maize and related grasses. Nature. 436, 1119–1126 (2005).

73. E. Bortiri, G. Chuck, E. Vollbrecht, T. Rocheford, R. Martienssen, S. Hake, ramosa2 encodes a LATERAL ORGAN BOUNDARY domain protein that determines the fate of stem cells in branch meristems of maize. Plant Cell. 18, 574–585 (2006).

74. N. Satoh-Nagasawa, N. Nagasawa, S. Malcomber, H. Sakai, D. Jackson, A trehalose metabolic enzyme controls inflorescence architecture in maize. Nature. 441, 227–230 (2006).

75. P. Bommert, C. Whipple, Grass inflorescence architecture and meristem determinacy. Semin. Cell Dev. Biol. 79, 37–47 (2018).

76. M. J. Scanlon, K. Ohtsu, M. C. P. Timmermans, P. S. Schnable, Laser microdissection-mediated isolation and in vitro transcriptional amplification of plant RNA in Curr. Protoc. Mol. Biol., (John Wiley and Sons Inc, 2009), chapter 25:Unit 25A.3. doi: 10.1002/0471142727.mb25a03s87.

77. Z. Dong, Y. Xiao, R. Govindarajulu, R. Feil, M. L. Siddoway, T. Nielsen, J. E. Lunn, J. Hawkins, C. Whipple, G. Chuck, The regulatory landscape of a core maize domestication module controlling bud dormancy and growth repression. Nat. Commun. 10, 3810 (2019).

78. L. Conti, D. Bradley, TERMINAL FLOWER1 is a mobile signal controlling Arabidopsis architecture. Plant Cell. 19, 767–778 (2007).

79. M. Can, W. Wei, H. Zi, M. Bai, Y. Liu, D. Gao, D. Tu, Y. Bao, L. Wang, S. Chen, X. Zhao, G. Qu, Genome sequence of Kobresia littledalei, the first chromosome-level genome in the family Cyperaceae. Scientific Data. 7, 1–8 (2020).

